# The key role of oxidation state and flake size in the ecotoxicological impact of industrial graphene oxide nanoflakes

**DOI:** 10.64898/2026.11.12.766197

**Authors:** Ahmed Subrati, Paolo Bigini, Ada De Luigi, Carmina Natale, Kahina Mehennaoui, Marianne Houin, Aline Chary, Tommaso Serchi, Tom Venus, Asia Saorin, Marco P. Monopoli, Xueqing Zhang, Kefei Zhao, Beatriz Alonso, Irina Estrela-Lopis, Changyou Gao, Alex Zabeo, Danail Hristozov, Sergio Moya, Luisa Diomede

## Abstract

Graphene oxide (GO) is a widely used nanomaterial across various scientific and technological fields. Environmental exposure to GO and consequent ecotoxicological effects have been largely reported. Despite the significant efforts and illuminating conclusions, establishing fair pairwise correlations between material and toxicity descriptors remains challenging and controversial. Key drawbacks in GO ecotoxicological studies to date include a lack of standardized materials, poorly defined descriptors, and insufficient material variants. Here, we establish the first comprehensive framework to tackle the multidimensional ecotoxicology of GO. We perform thorough material characterization to derive descriptors that precisely define four key GO variants used in industrial scale-up, with differing flake granularity and oxidation extent. These GO variants are evaluated against toxicological endpoint descriptors across two animal models (*Daphnia magna* and *Caenorhabditis elegans*) and two plant models (rice and pepper). Our findings underscore the inherently multidimensional nature of GO toxicity by identifying key physicochemical properties associated with it. This study provides an unprecedented platform for parametrizing GO materials using material and toxicity descriptors to contribute to nanomaterial safety.

## Introduction

Our society is witnessing ever-increasing development and use of nanoscale materials across multiple sectors: manufacturing, household, agriculture, medicine, energy, and others. Among these materials, graphene oxide (GO) nanoparticles (NPs) are increasingly playing a key role across multiple areas due to their unique physicochemical properties arising from their layered structure and sp^2^ carbon arrangements. Structurally, GO can be described as a derivative of pristine, honeycomb-structured sp^2^ carbon graphene. It is functionalized on both its basal planes and edges with oxygen-containing groups, including epoxides, hydroxyls, ethers, carbonyls, lactones, lactols, and carboxyls [1, 2]. The large and increasing use of GO, its tendency to associate, and its low degradability raise potential ecotoxicological concerns, particularly regarding the possible acute and longer-lasting effects resulting from exposure of animals and plants. Compared with other carbon-based materials, such as carbon nanotubes, there is no evidence to date that associates GO with an increased carcinogenic risk [3]. However, a series of studies has reported potential cytotoxic effects of certain types of GO nanoflakes [4–6]. With respect to human and animal health, several points, such as the generation of superoxide species and their ability to cross biological barriers or be absorbed into the circulation after ingestion or inhalation, are of particular interest.

Toxicological endpoints of GO have been evaluated in eukaryotic cells and rodents for several years [7, 8]. In zebrafish, GO exposure significantly increased malondialdehyde, catalase (CAT), and superoxide dismutase (SOD) levels, accompanied by a reduction in glutathione content in vacuolated liver cells, demonstrating a pro-oxidant effect in vivo. In environmental science, it has been reported that GO alters the functional expression of certain pollens, reducing the fertilization capacity of certain seed plants and causing severe fertilization problems.

Despite the emerging body of studies on GO nanosafety, comparisons across various animal and plant biological matrices are difficult due to heterogeneity in exposure types, concentrations, and intended outcomes [9]. There is currently a lack of rigorous studies that test GO materials, specifically those produced for industrial use, across diverse biological matrices, including fluids, cells, small animals, and plants. In this context, a correct assessment of the overall risk must focus on comparing GO nanoflakes and nanosheets with different chemical characteristics [10]. In fact, GOs with different flake sizes and oxidation levels can differ in important parameters, including degree of stacking, polydispersity index, stability in media, and interactions with plasma proteins or environmental fluids [11, 12].

In this study, we aim to correlate GO physicochemical properties with its toxicity in both animal and plant models. To enable a comparative analysis, four types of GO nanoflakes, currently used in industrial-scale-up, and characterized by different oxidation states and flake sizes, were selected.

We will first analyze how these two characteristics affect other properties of the GOs, mainly aggregation and stacking degree, aggregate size, and their interactions with biomolecules, which will be determinants of toxicity. Toxicological endpoints were then measured in two simplified animal models, namely the crustacean *Daphnia magna* and the nematode *Caenorhabditis (C.) elegans*, which are widely used for nanosafety testing, as well as different types of plants, seeds, and vegetal cells. Our findings underscore the inherently multidimensional nature of GO toxicity, first identifying key physicochemical properties relevant to it and offering an unprecedented platform for parameterizing GO materials using material and toxicity descriptors to contribute to nanomaterial safety.

## Results and discussion

### Physicochemical characterization of GOs

The characterization of the four synthesized GOs was performed by determining key physicochemical features, including size, shape, and aggregation state. Scanning electron microscopy (SEM) analysis revealed sheer morphological differences among the four GO materials. GO1 and GO4 particles shared analogous topography, characterized by smooth, wrinkle-poor surfaces (**Fig. 1** and **Table S1**). These two materials exhibited similar degrees of oxidation, as evidenced by the oxygen weight percentage measured by elemental analysis. GO2 particles with the lowest oxidation extent exhibited numerous defects, folds, and ripples, as inferred from the sharp contrast. Such defects were caused by the destructive reduction process that restores the graphitic carbon backbone (sp^3^_→_sp^2^), triggering immense stress and deformation that cause defects and particles to crumple onto themselves. GO3 particles, with a medium degree of oxidation, exhibited morphological features ranging from GO1/GO4 to GO2, with smooth surfaces and discernible, contrasting folds and ripples. Particle size, approximated as the circular equivalent diameter, is determined following number-weighted and area-weighted distributions. Using the cumulative distribution parameter D50, it is possible to determine the order of GO sizes: (largest) GO1 > GO4 > GO3 > GO2 (smallest). Both distributions reflect the true granularity of GO; the number-weighted distribution highlights the significant presence of sub-micron particles, often featureless, whereas the area-weighted distribution highlights the exemplary flake particles with discernible features. Transmission electron microscopy (TEM) analysis furthermore corroborated the SEM data. On the one hand, images from GO1 and GO4 were transparent due to minimal folds and ripples. On the other hand, GO2 particles appeared less transparent due to densely packed defects and ripples. GO3 particles were characterized by medium transparency, indicative of a moderate density of defects, folds, and ripples. Morphology and thickness of GO particles were also analyzed using Atomic Force Microscopy (AFM), indicating that GO2 particles exhibited the widest interlayer stacking, followed by GO1, GO3, and GO4. This ordering reflected the nature of GO particles; GO2 was the least oxidized, whereas GO4 was the most oxidized, and the exfoliated derivative of GO1. Analysis of C 1s XPS reveals distinct chemical environments in the GO materials. A major descriptor was the carbon-to-oxygen atomic ratio (C/O), which indicates GO hydrophilicity; a value closer to unity indicates a higher degree of oxidation and greater dispersibility in aqueous media. The C/O values for GO materials are ordered in ascending order: GO4 _≤_ GO1 < GO3 < GO2 (**Table S2**). The presence of sulfur can be traced to the oxidation of GO1, in which concentrated sulfuric acid was used, and the residual sulfur content may appear as covalent sulfates [19]. The nitrogen present in all samples can be attributed to impurities in the source graphite used to synthesize GO1, from which the other GO materials are derived (**Table S2**). Local chemical environments were probed via peak fitting of deconvolved C 1s XPS (**Fig. 1, Table S3**). Various chemical environments characterize different carbon atoms: C=C in graphitic domains (284.75 eV), C C in amorphous domains (285.29 eV), C O in hydroxyl/epoxy (286.79 eV), C=O in carbonyl/quinone (288.03 eV), COO in carboxyl/lactol/lactone (289.05 eV), _π π_* transitions/interactions (290.95 eV) [13, 20, 21]. While GO1 and GO4 shared a similar bimodal profile in their C 1s XPS spectra, GO2 showed a unimodal profile. The GO3 profile was bimodal, with the C O component relatively less intense than in GO1/GO4. The C/O ordering was consistent with the relative abundance of the local chemical environments of carbon atoms. In particular, the restoration of graphitic C=C bonds can be observed in GO2 and GO3, the most- and mildly reduced GO derivatives of GO1, respectively. The C-O component also varied considerably across the GO materials; it was lowest in GO2 and highest in GO1/GO4, with the C-O concentration in GO3 falling between these two extremes. Survey XPS proved the absence of impurities in all GOs, as shown in **Figure 2A**.

**Figure 1.**
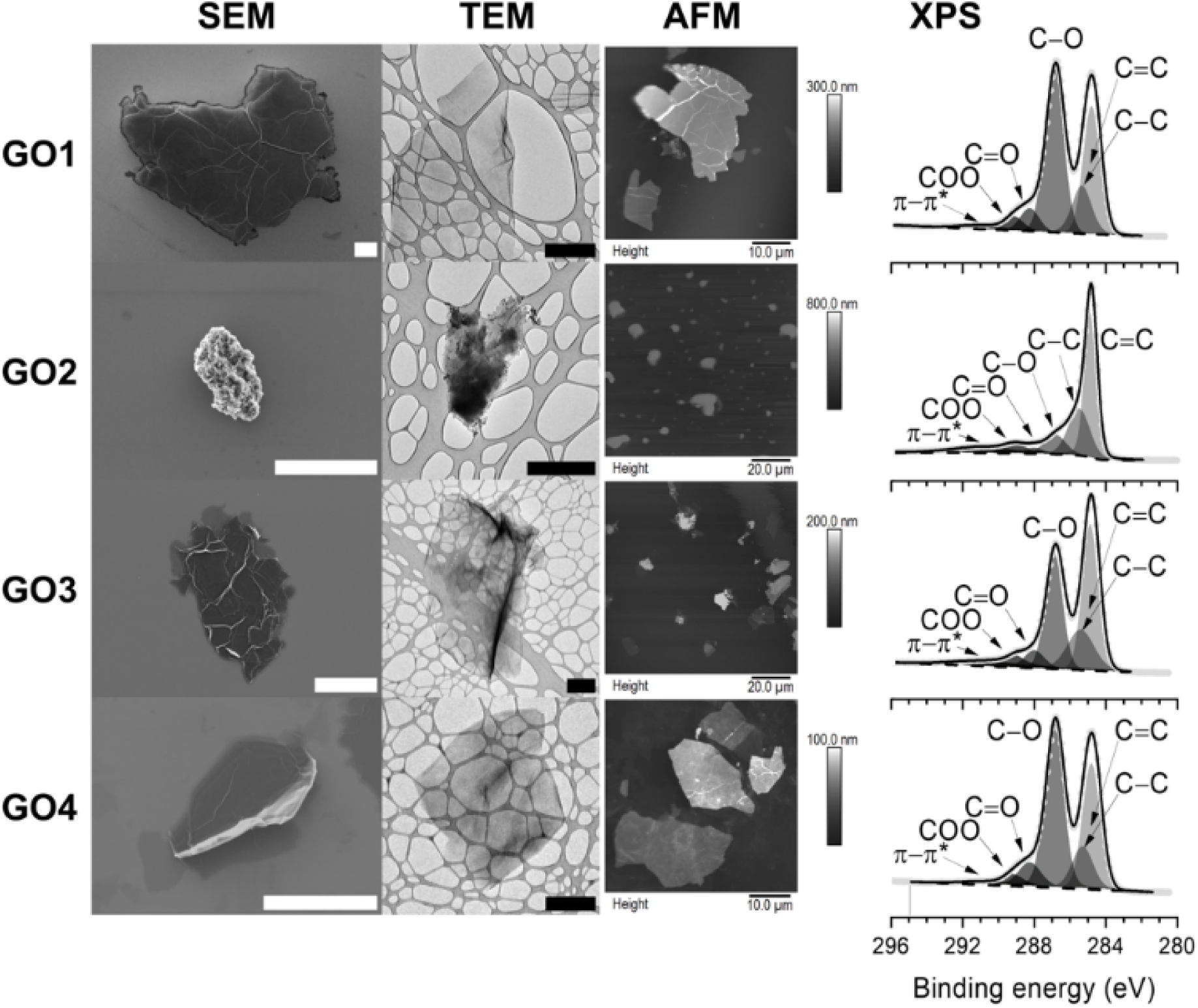
Physicochemical characterization of GOs. Representative images of SEM (scale bar: 5 μm), TEM (scale bar: 2 μm), and AFM analyses, and C 1s XPS obtained for the different GOs. Three microscopic methods and one spectroscopic method reveal the distinct differences among the GOs. The EM images, in particular, reveal the particle electron density and beam-interaction effects when probing fine features of observed particles. The thick, electron-dense GO1 flake interacts sufficiently with the electron beam, thereby illuminating extensive ripple and fold features. The highly crumpled and defective GO2 particle has an enormous electron density, which causes it to appear partly opaque in TEM and to appear clearly with sharply contrasted edges in SEM. GO4 exhibits less intense surface features than GO1 due to its overall lower electron density. The GO3 particle combines the sharp, contrasting features of GO2 with the folded and rippled features of GO1/GO4. The bimodal C 1s XPS signal is a fingerprint of GO. Cumulative contributions of various chemical carbon environments yield significant shoulders and features in the high-resolution C 1s XPS of GO

**Figure 2.**
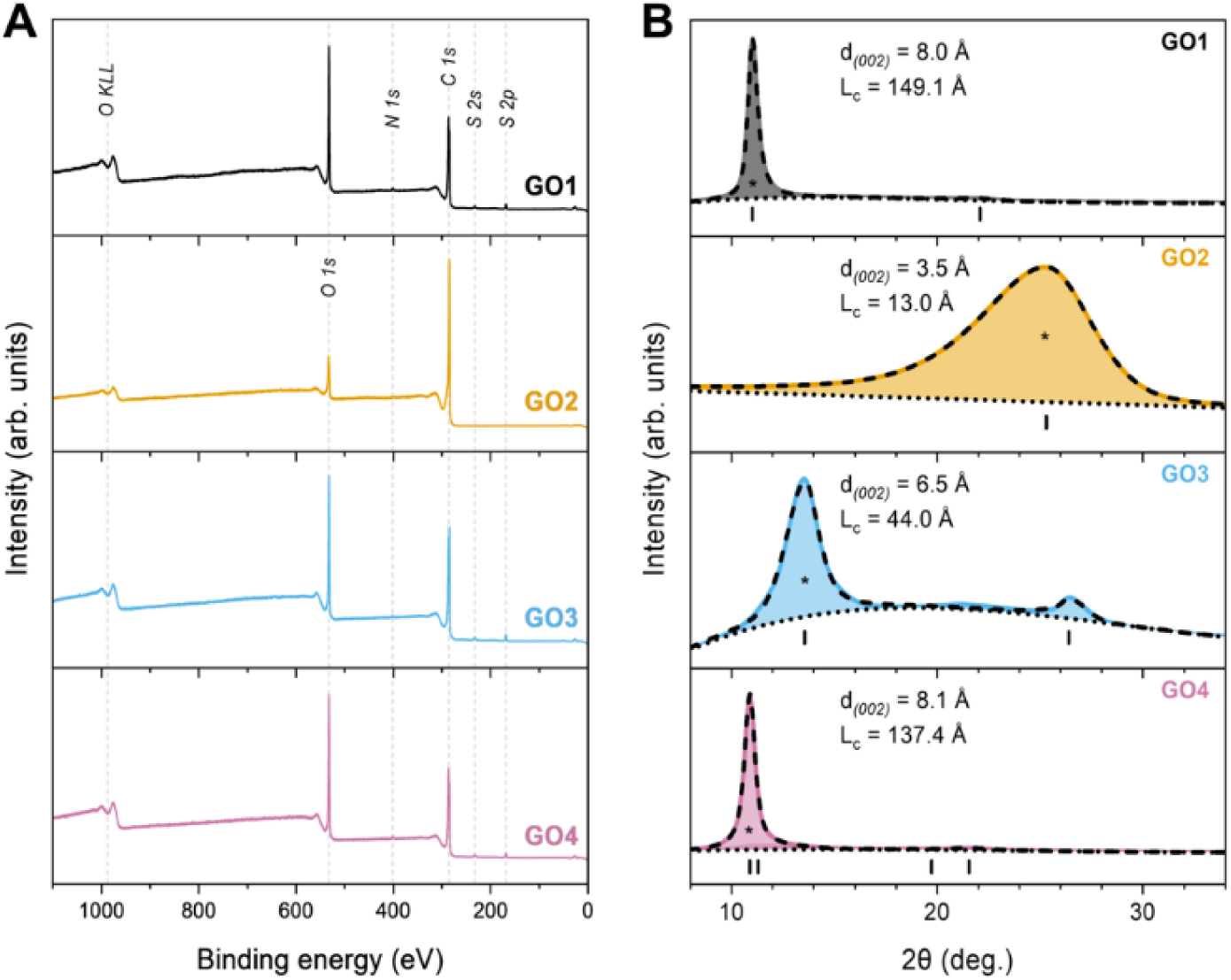
XPS composition and X-ray diffractograms analysis. (A) Wide-scan XPS survey and (B) X-ray diffractograms (XRD) of GO1, GO2, GO3, and GO4. Notice the changes in the 2θ position and width of the primary peak (indicated with an asterisk), which are closely associated with the chemistry of GO. Thick color line: raw XRD pattern; dashed line: cumulative fit; dotted line: background; color-integrated area: fit component; and short vertical line: position of the Bragg reflection. Interlayer spacing (d) and crystallite c thickness (Lc) values are reported for each GO.

The distinct chemistry of GO materials affected interlayer interactions and interlayer spacing. X-ray diffractograms, shown in **Figure 2B**, indicate that GO1 and GO4 exhibited signals at 2_θ_ of 11.01° and 10.89°, corresponding to interlayer spacings of 8.0 Å and 8.1 Å, respectively. A signal at 2_θ_ of 25.30° was observed in the X-ray diffraction (XRD) pattern of GO2 corresponding to an interlayer spacing of 3.5 Å. The XRD pattern of GO3 showed two signals at 2_θ_ values of 13.56 ° and 26.40 °, corresponding to interlayer spacings of 6.5 Å and 3.4 Å, respectively. The secondary XRD signal of GO3 indicated the retention of graphitic crystals, while the primary peak indicated the presence of sterically hindering oxygen functional groups within the interlayer domains; however, with a lower abundance relative to GO1/GO4 [2, 22]. This directly reflects the impact of the lower mass equivalents of KMnO_4_ used in GO3 synthesis. The most spacious interlayer spacings of 8.0 Å and 8.1 Å in GO1 and GO4 were a direct consequence of the highest oxidation extent. GO2 had the narrowest interlayer domains, indicating the lowest abundance of oxygen functional groups. Another significant feature was the width (full width at half-maximum, FWHM) of the diffraction signals. The primary peaks for GO1, GO2, GO3, and GO4 had FWHM values of 0.56°, 6.52°, 1.90°, and 0.61°. This indicates the presence of the thinnest crystals in GO2 and the thickest crystals in GO1. In GO2, restored sp^2^ domains exhibited volume expansion and protrusion of amorphous sp^3^ domains, which render smaller crystals [23]. Notice how the GO3 FWHM of the primary XRD peak was larger than that of GO1/GO4 and smaller than that of GO2, corroborating that GO3 is the GO1 derivative with a medium oxidation extent. The effect of sonication was evident in the slight increase in the FWHM of GO4 compared to GO1.

The physicochemical properties of GOs were also characterized using label-free high-resolution confocal Raman microscopy (CRM). Bright-Field (BF) stitched imaging, covering areas up to several mm^2^, was combined with Raman imaging (RI) to visualize particle morphology and to enhance detection even beyond the optical resolution limit, enabled by the high Raman activity of the D and G bands. The combined RI and BF image, in which each identified particle was chemically verified by its corresponding Raman spectrum, was analyzed using a particle-searching algorithm to locate, identify, categorize, and characterize individual particles. CRM images revealed distinct morphologies among the four GO materials (**Fig. 3A**).

**Figure 3.**
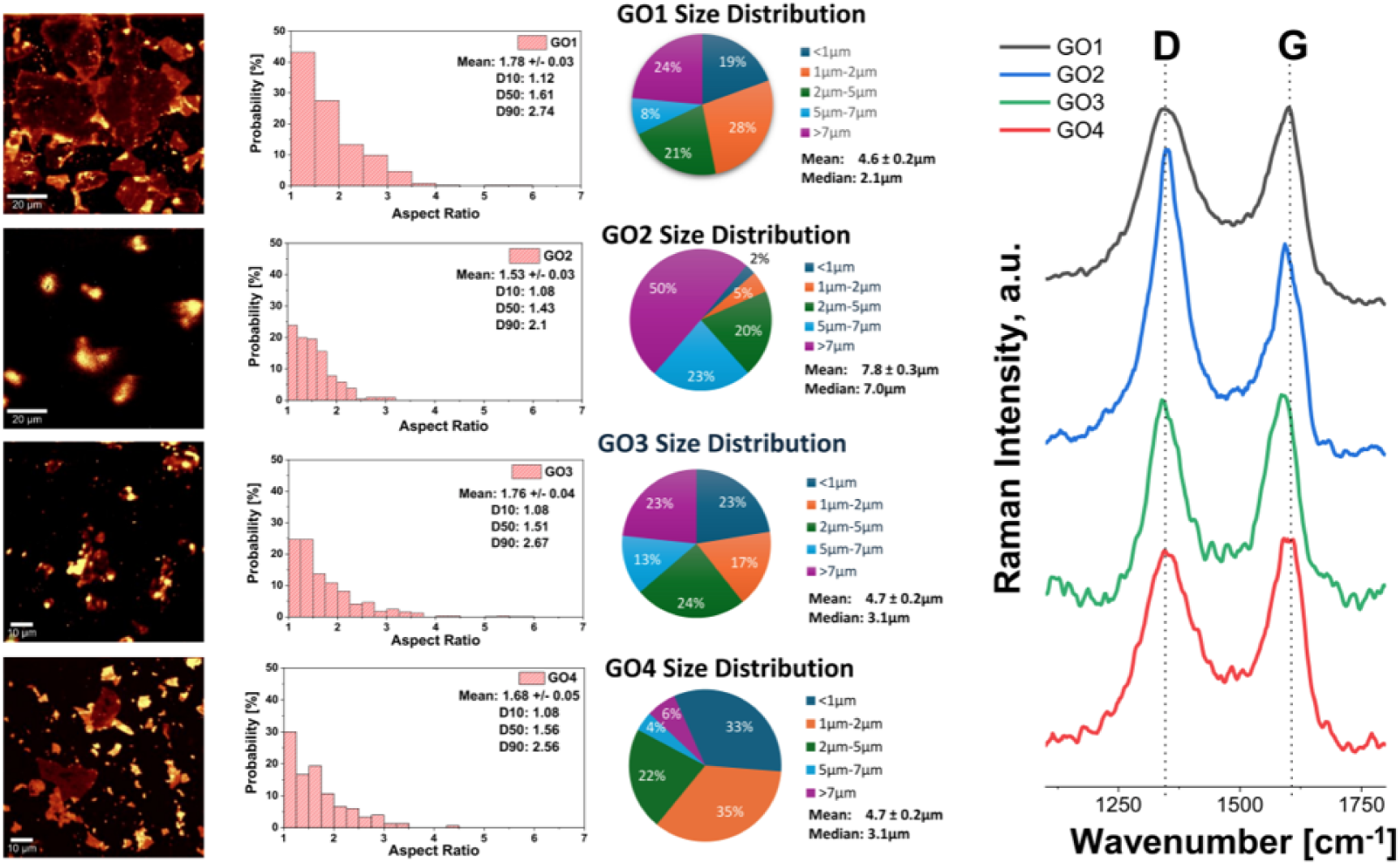
CRM characterization of GO particles. (A) **High-resolution** confocal Raman images; (B) aspect ratio distribution of individual particles; (C) size composition shown as pie charts; (D) averaged Raman spectra extracted from the corresponding Ris.

The relative thickness of the flakes was determined by calculating the integrated intensities of the D and G bands, which were visualized in color-coded Raman images. Particles of GO1 and GO4 with high oxidation levels primarily exhibit smooth, thin flakes with very low Raman signal intensity. A significant number of small GO particles, down to 250 nm in size, were notably absorbed on the surface of larger, transparent (dark brown) GO1 flakes, which exhibited relatively few folds. In contrast, the reduced GO2 and less-oxidized GO3 samples exhibited more three-dimensional structures, with flake thicknesses up to 2 µm in GO2. These observations are consistent with SEM, TEM, and AFM already described in Figure 1. Approximately 2000 individual particles were analyzed and characterized at different positions on the filter. The aspect ratio (AR) followed a log-normal distribution for all GO samples, ranging from a ratio of 1 (circular shape) to 4 (elongated flake) (**Fig. 3B**). Particles with a circle-equivalent diameter of less than 1 µm were predominantly circular. Up to 30% of GO1 and GO4 particles exhibited laterally elongated shapes (2>AR> 4). On the other hand, only 12% of GO2 particles displayed elongated morphology. The percentile particle values of AR for the studied GOs correlate well with the SEM results (**Table S1**).

The size distributions of the different GO samples were highly polydisperse, spanning the nano-to microscale (**Fig. 3C**). The size distributions of GO1, GO4, and GO3 exhibit strongly asymmetrical long-tailed distributions, with significant differences between the mean and median, whereas GO2 follows an almost Gaussian distribution. The median values of GO1, GO4, and GO3 particles are shifted strongly towards the smaller sizes compared with the number-weighted values presented in **Table S1**. This difference can be explained by the improved detection of particles ranging from 280 nm to 1 µm using high-resolution RI. Furthermore, a large number of particles smaller than 1 µm can be distinguished and analysed on the surface of larger particles as a result of the drying process (**Fig. 3A**, upper), enabled by the enhanced image processing. Around 19%, 33%, and 23% of the analysed GO1, GO4, and GO3 particles were below 1 µm in size, respectively. By comparison, only 2% of GO2 were smaller than 1 µm. The largest mean CE diameter was observed for GO2 particles, which have a strong tendency to form aggregates in aqueous solutions.

Considering toxicologically relevant particles smaller than about 2 µm [24, 25], approximately 47% of GO1, 68% of GO4, 40% of GO3, and only 7% of GO2 particles fall within the size range capable of crossing the cell membrane and being internalized. **Figure 3D** shows a strong blue shift of about 10 cm-1 in the G-band position for GO1 and GO4. This shift was observed for both types, indicating a high degree of oxidation [26]. In the case of reduced GO2 and less-oxidized GO3, a redshift of the G band was observed, a phenomenon known as “graphitic self-healing”. These results are in good agreement with the XPS data, which show the highest oxygen content in the GO1 and GO4 samples (**Table S2**). Analysis of Raman spectra of GO materials (**Fig. 4A**) can provide insight into the chemical and physical changes in GO sheets and the probable transformations affecting the GO carbon backbone. The Raman spectra were deconvoluted into 9 bands: D*, D, D”, G, D’, M, 2D, D+D’, and 2D’ (**Table S4**) and, the peak integrated intensity ratio between the D band and the G band (AD/AG) was employed as an excellent descriptor of defect density, from which one can estimate the crystallite width (La). Interestingly, the peak integrated intensity ratio between the D band and the D’ band (AD/AD’) can help identify the nature of defects [27], the highest defect density and oxidation extent in both samples (**Table S5**). The lowest AD/AG value was observed in GO2, indicating the fewest defects induced by oxygen functionalities and the highest restored C=C, a finding supported by XPS. The defects in GO3 were representative of a medium oxidation level, lying between GO1/GO4 and GO2. The AD/AD’ values for GO1 and GO4 were associated with vacancy-type defects, whereas the values for GO2 and GO3 were associated with sp3-type defects (**Fig. 4B**).

**Figure 4.**
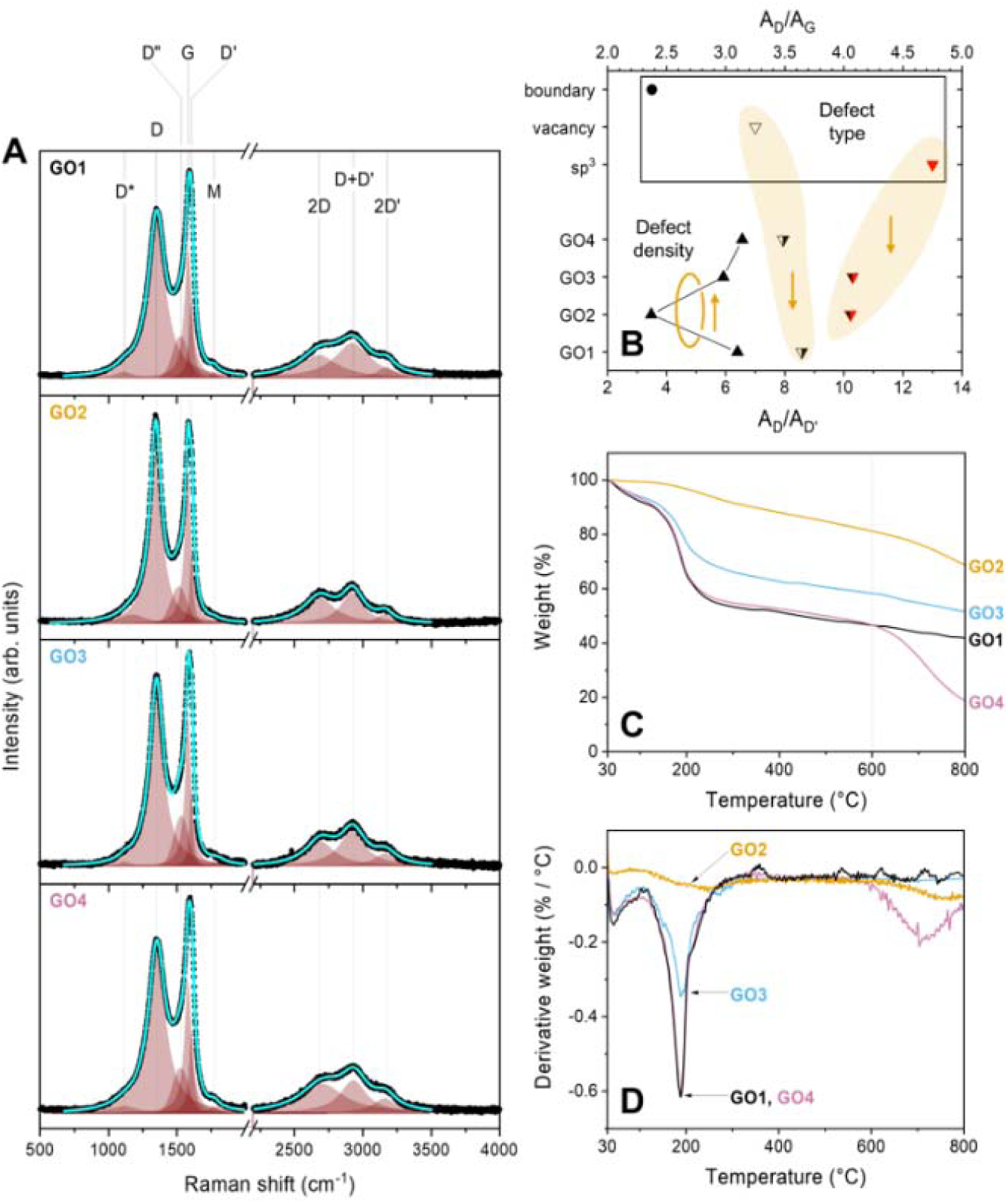
Raman spectra, thermograms, and derivative thermogravimetry (DTG) curves. (A) Analysis of the Raman spectra of GO1, GO2, GO3, and GO4 reveals the contributions of nine distinct components. Black scatter points: experimental Raman spectra; cyan line: cumulative fit; color-integrated area: fit component. (B) Descriptors of defect type (AD/AD’, downward-pointing triangles) and density (AD/AG, upward-pointing triangles) correlation via Raman spectroscopy analysis. (C) Thermograms and (D) DTG curves of GO1, GO2, GO3, and GO4.

TGA provided a descriptor of the oxidation extent of GO materials; the higher the oxidation extent of GO (C/O closer to unity), the bigger the mass loss associated with the oxygen-functionalized proportion of the sample. Scrutiny of thermograms in **Fig. 4C** indicated that all GOs possess distinct profiles. GO2 exhibits the smallest oxygen-functional mass loss, as there were no clear steep curves. This is reasonable given the overall low oxidation extent of GO2 (C/O = 9.16). GO1 and GO4 shared a similar profile in their respective thermograms, with slight variations attributable to the average smaller, thinner flakes in GO4 compared to GO1. The GO3 thermogram profile lies between those of GO1/GO4 and GO2, indicative of a medium oxidation extent. In attempts to obtain numerical descriptors from profiles, it was found that the thermal degradation reaches a distinct point around 600 °C. At that temperature, GO1 and GO4 thermograms intersected. This is inevitable since GO4 is derived from GO1 and hence possesses a similar degree of oxidation. For this reason, and in light of other indications from the literature, it is recommended to treat this as an indirect descriptor of oxygen-functionality mass loss in GO materials: 100-m600, where m600 is the percent mass at 600 °C, as read from the thermogram [2]. To validate such a descriptor, it is therefore quite intuitive to associate its display with the experimental acquisition conditions (as mentioned above). Based on the 100-m600 descriptor for GO materials, GO1, GO2, GO3, and GO4 have respective oxygen-functionality mass losses of 53.4%, 18.8%, 41.8%, and 53.4%. As depicted in **Fig. 4D**, the DTG curves revealed distinct stages during which the GO carbon backbone undergoes rapid thermal decomposition into COx and H_2_O. All GO samples exhibited a common stage of mass loss centered at approximately 188 °C, attributed to the thermal decomposition of GO oxygen functionalities to COx and H_2_O. It is also significant to highlight the plausibility of CO-induced self-reduction mechanisms, in which freshly released CO from the GO carbon backbone reacts with neighbouring oxygen functionalities to generate CO_2_ (and H_2_O), thereby further increasing the reduction extent [28, 29]. The main parameters of the four different GOs synthesized for this study are summarized in **Table S6**.

### Impact of GOs on *C. elegans* and *D. Magna*

It is widely known that the ability of nanomaterials to interact with biological targets is dramatically influenced by the medium. To investigate this point, we used Differential Centrifugal Sedimentation (DCS) to assess the sedimentation of GOs in a sucrose gradient before and after exposure to biological fluid. While the DCS size measurements are not accurate for nonspherical objects, this technique allows differentiation between agglomerates and clusters of different sizes and densities, as they will sediment through the gradients separately. Additionally, the DCS is a reliable and reproducible technique that provides relative shifts of the same sample after surface modification or corona formation [30–32]. Before evaluating the impact of GOs on *C. elegans* toxicity, we examined the behavior of the different materials in the biological media used for worm experiments: water used to wash a solid Nematode Growth Medium (NGM) plate containing OP50 *E. coli* as food, and water alone as a negative control. The DCS analysis showed notable differences in the GO1-GO4 after their exposure to water. GO1 and GO3 had a similar size distribution, where the materials were distributed between two peaks with the apparent size distribution below 1 and 2 nm respectively, where smaller flakes had a predominant presence, while GO2 and GO4 displayed broader distribution (**Fig. 5**). After a 1-hour incubation, DCS analysis revealed a general similarity between the profiles of GO dispersions in water alone and in water from NGM plates with or without *E. coli* (**Fig. 5**).

**Figure 5.**
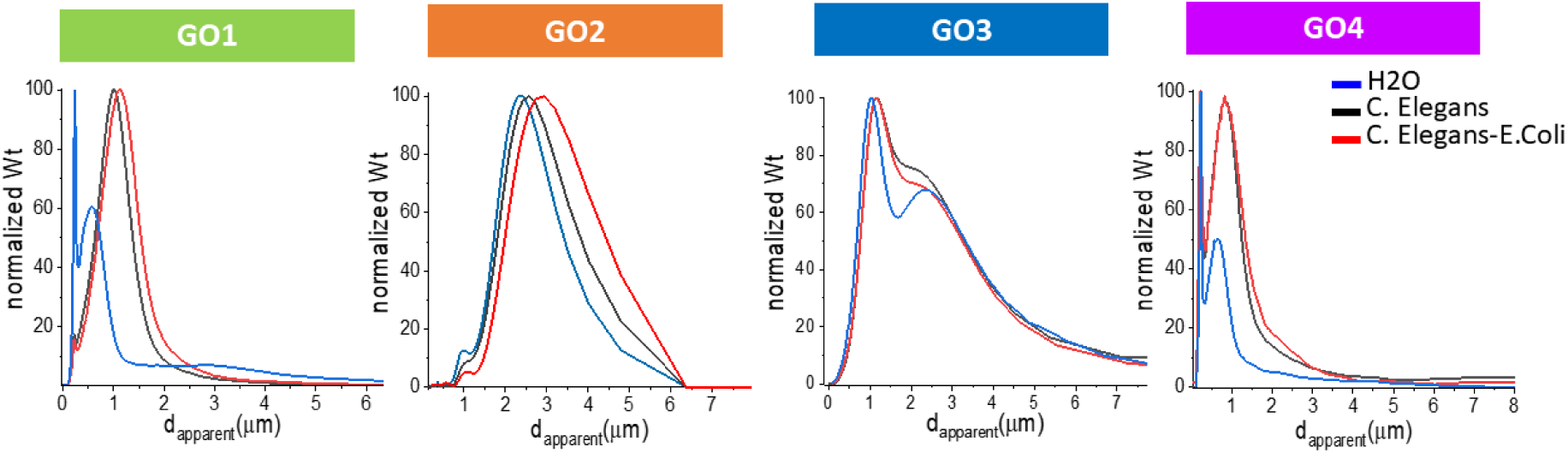
DCS analysis of GOs dispersed in biological fluids used in *C. elegans* experiments. GO size distribution comparison after exposure in water (control), *C. elegans* culture media with water (*C. elegans* + H2O) or *C. elegans* culture media with OP50 *E. coli* (*C. elegans* + *E. coli*). The GOs were exposed to the media for 1 hour at 37 °C.

The difference between GO4 and GO1 was significant: the peak at small size almost completely disappeared in GO1, whereas it remained in the GO4 dispersion (**Fig. 5**), indicating that the small flakes and GO complexes are less stable in the presence of complex fluid and form aggregates, while a persistent size distribution was observed for GO2 and GO4 after exposure to the environmental fluids.

The acute effect of GOs on *C. elegans*’ viability was then investigated by treating worms with different concentrations (0.06-1 mg/ml) of materials. **Figure 6A** shows a dose-dependent decrease in worm viability 24 hours after administration of GO1, GO3, and GO4 but not GO2. Among the various GOs, GO4 was the most toxic, with an IC50 value of 0.14 mg/ml, half that of GO3 (0.29 mg/ml) and one-third that of GO1 (0.42 mg/ml). After 24 hours of treatment at the highest dose of 1 mg/ml, GO1, GO3, and GO4 similarly reduced worm viability, with viable-worm percentages of 48.4%, 48.5%, and 47.9%, respectively, and toxicity did not increase further over time (**Fig. S3A**). The data obtained from measuring the lifespan and the health span of worms confirm that GO1, GO3, and GO4 can also exert a subtoxic effect (**Fig. 6B, Table S7**). At 0.125 mg/ml, GO1 and GO3 similarly reduced the worms’ lifespan by less than 10% relative to vehicle-fed worms, whereas GO4 significantly shortened it by approximately 23%. Additionally, GO4 impaired the worms’ spontaneous movement, resulting in a significant 20% reduction in median health span (**Table S7**).

**Figure 6.**
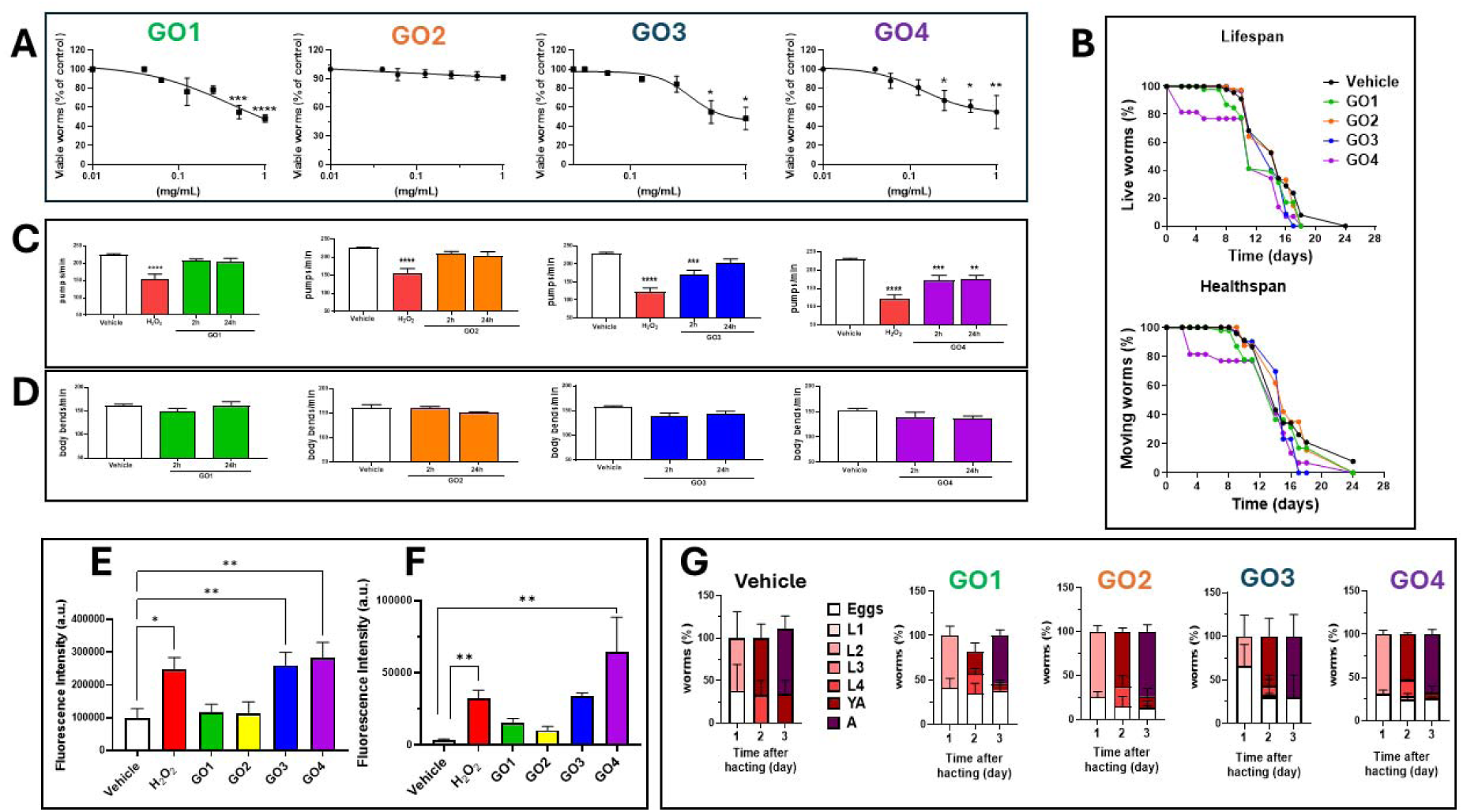
Acute and subacute toxic effects of GOs on *C. elegans*. **A**) Synchronized worms were fed 0.06-1 mg/ml different GOs suspended in water without OP50 *E. coli* (50 worms/50 µl). Worms were fed water alone (Vehicle) (50 worms/50 µl) as a negative control, and 10 mM H2O2 as a positive control (50 worms/50 µl). After 2 h of incubation at 20°C, worms were plated on NGM plates seeded with fresh bacteria, and the number of viable worms was evaluated 24 h later. Data are the mean of viable worms (% of vehicle) ± SEM (N=50) from 3 independent experiments. *p< 0.05, **p<0.01, ***p<0,005, and ****p<0.001 *vs.* Vehicle, one-way ANOVA and Bonferroni’s *post hoc* test. **B**) Lifespan and health span curves of worms treated as described above with 0.125 mg/ml GOs or water as control (Vehicle) (50 worms/50 µl). Dead, alive, and censored animals were scored. Data are the mean ± SEM (N=50) from 3 independent experiments. The median lifespan, health span, and statistical analyses are shown in Table S7. **C**) Pharyngeal pumping and **D**) locomotor activity of worms treated as described above with 0.125 mg/ml of different GOs suspended in H2O (50 worms/50 µl). Control worms were fed the same volume of H2O alone (Vehicle), and positive control worms were fed 1 mM H2O2. Pharyngeal pumping (pumps/min) and locomotor activity (body bends/min) were scored 2 and 24 h after plating worms on NGM plates seeded with OP50 *E. Coli*. Data are the mean ± SEM (N = 30) from 3 independent experiments. ** p< 0.05, ***p<0.001, and ****p < 0.0001, one-way ANOVA and Bonferroni’s *post hoc* test. **E, F**) Relative quantification of MitoSOX™ Red fluorescence intensity measured with Fiji imaging software in the **E)** whole body and **F**) pharynx of worms. Data are the mean ± SEM (N = 12). * p<0.05 and ** p<0.005, one-way ANOVA, and Bonferroni’s *post hoc* test. **G**) Effect of GOs on the development of nematodes. Worms (50 worms/50 µl) were fed for 2 hours with 0.125 mg/ml GOs suspended in water, without OP50 *E. coli*, then plated on NGM plates seeded with the bacteria. Control worms were fed with only water (vehicle). The percentages of eggs and worms at the L1, L2, L3, and L4 larval stages, and the young-adult (YA) and adult (A) stages, were determined at 1, 2, and 3 days after egg hatching. Data are the mean ± SEM (N = 12).

The subtoxic effects of GOs were also evaluated using behavioral trials, including assessments of feeding, locomotion, reproduction, and development. First, the feeding behavior and motility 2 and 24 hours after administering 0.125 mg/ml GOs were considered (**Fig. 6C, D**). GO1 and GO2 did not affect the worms’ pharyngeal or motility functions under any tested conditions. GO3 led to a transient 25% impairment of pharyngeal contraction, statistically significant at 2 hours but not at 24 hours after administration, whereas GO4 caused a permanent 25% impairment of pharyngeal contraction. Hydrogen peroxide, used as a positive control, reduced the pharyngeal contraction by 47%. No alteration in worms’ motility was indeed observed by both GO3 and GO4. These results are consistent with those from the acute toxicity study and indicate that GO4 was the most impactful in *C. elegans* among the GOs.

It has recently been reported that GO can act as an environmental oxidative stressor in worms, disrupting cellular functions and leading to irreversible DNA damage [33]. Thus, we investigated whether the effects of the GOs on pharyngeal contractions were related to their mitochondrial superoxide, evaluated by MitoSOX™ Red staining.

The fluorescent signals from whole worms and the pharynx (**Fig. S4**) were quantified, and the data are shown in **Figures 6E** and **F**. As expected, the treatment with 1 mM H2O2 led to a significant increase in the signal in both the whole body of worms and the pharynx. No increase was observed in worms treated with 0.125 mg/ml GO1 and GO2 compared to the vehicle-treated group. In worms treated with 0.125 mg/ml GO3 and GO4, we observed a significant increase in fluorescence in the whole body, comparable to that caused by H2O2 (**Fig. 6E**). A similar increase was also observed in the pharynx, although significant for GO4 only (**Fig. 6F**). These results suggest that the higher acute and subacute toxicity recorded in animals exposed to GO4 is related to its ability to induce mitochondrial superoxide production. At 0.125 mg/ml, all types of GOs, including GO2, significantly affected worm development (**Fig. 6G**), mainly delaying egg hatching compared to vehicle-treated worms. In fact, unlike vehicle-fed worms, eggs were still present in worms treated with GOs 2 and 3 days after hatching, significantly reducing the number of young adult and adult worm generations. On the other hand, no effect on the worms’ overall reproductive capacity was found for all GOs (**Fig. S3B**). Overall, these data clearly indicate a hierarchical effect in *C. elegans*, with GO4 exhibiting the stronger toxic effect, whereas GO2 was apparently safe, except for delayed development.

To determine whether the differential effect of GOs could be extended to other animal models, the study was conducted in the crustacean *D. magna*. Interestingly, the impact of GO4 on acute toxicity was similar to that observed in *C. elegans* (*Table S7*). Based on the overlap of 95% confidence intervals, the relative toxicity ranking was: GO4 > GO1 _≥_ GO3 > GO2 (**Table S8**). The observed acute toxicity values align with those reported for other graphene-based materials, in which no immobilization was detected below 10 mg/l for graphene ink, and EC_50_ values ranged from 52 to 60 mg/l depending on the formulation [34].

As in nematode studies, heart rates were measured in *D. magna*. Average heart rates in the control group were 283.8 ± 8.5 and 258.1 ± 9.1 beats per minute (BPM) after 24 and 48 h of exposure, respectively. Significant decreases in heart rate were observed following GO4 exposure: 6% and 17% reductions at 7 and 78 mg/l after 24 h, respectively, and a 7% reduction at 78 mg/l after 48 h. However, recovery was observed when organisms exposed for 24 h were transferred to clean medium for 48 h, as heartbeat frequencies returned to levels comparable to those of controls (**Fig. 7**, upper panels). In addition, the distance procured by *D. magna* was significantly affected by GO4 exposure in a concentration-dependent manner (**Fig. 7**, lower panels). After 24 h of exposure, daphnids exposed to 0.7 mg/l showed no significant reduction in swimming activity compared to controls, whereas at concentrations of 7 and 78 mg/l, significant decreases of 32% and 41%, respectively, were observed.

**Figure 7.**
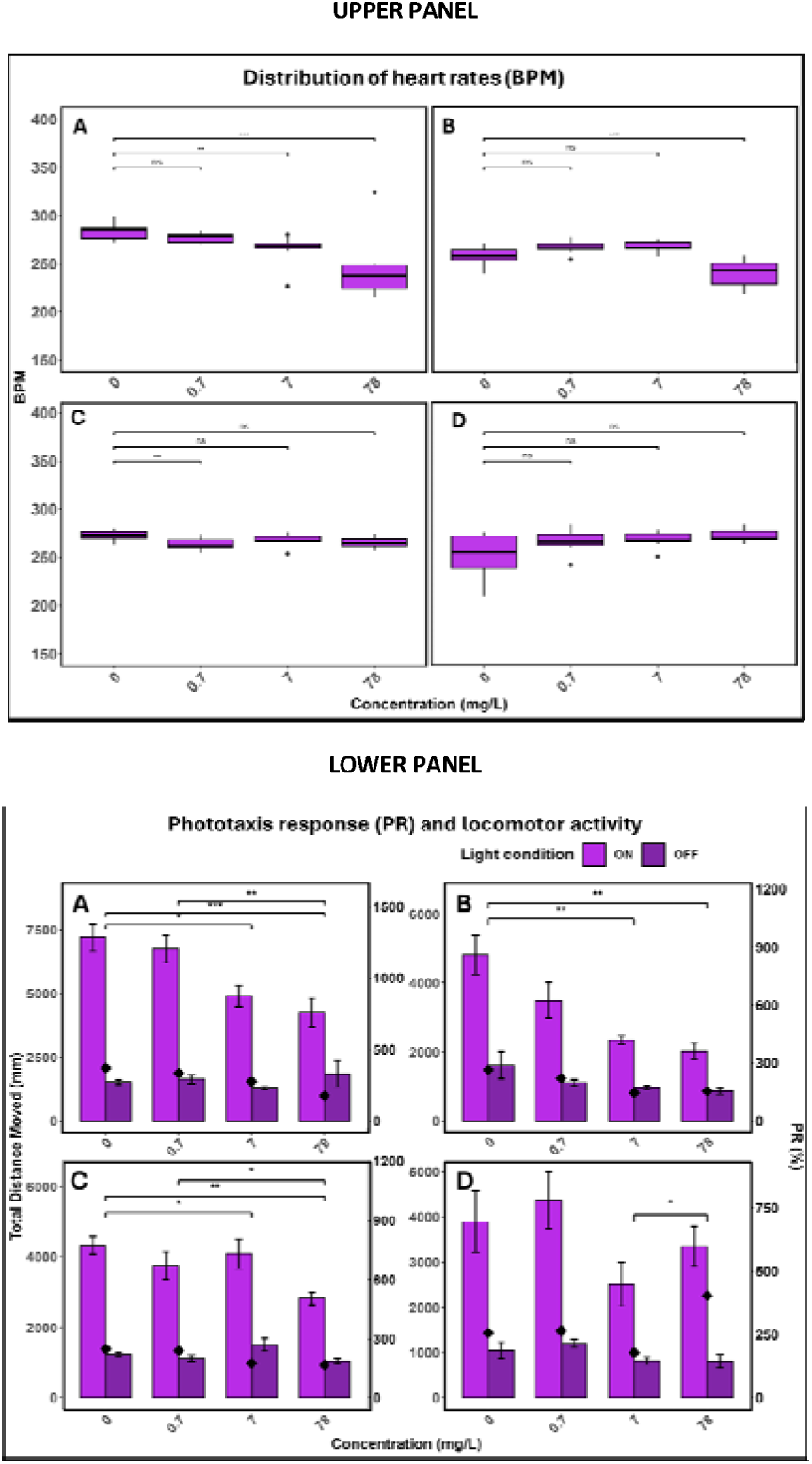
Impact of GO4 on *Daphnia Magna*. Distribution of heart rates, expressed as BPM, as a function of exposure concentration and experimental protocol. Boxes represent the interquartile range (IQR), the central line indicates the median, and individual points represent observations. Data are the mean ± SD. Brackets with asterisks indicate statistically significant differences between concentrations. *p<0.05, **p<0.01, ***p<0.001 according to Kruskal-Wallis and Dunn’s *post hoc* test. Each group corresponds to a different exposure/depuration protocol (A-D). Tested concentrations of GO4 were 0 (control), 0.7, 7, and 78 mg/l. Phototaxis response (PR) and locomotor activity under light ON/OFF conditions across exposure protocols and concentrations. PR was calculated as: PR (%) = ((DistanceON - DistanceOF F)/ (DistanceOFF)) × 100, where DistanceON represents the total distance traveled under light, and DistanceOFF represents the total distance traveled in the absence of light. Positive PR values indicate attraction to light, negative values indicate repulsion from light, and values close to zero indicate no significant response. Panels A-D correspond to the following protocols: A = 24 h exposure; B = 48 h exposure; C = 24 h exposure + 24 h depuration; and D = 48 h exposure + 48 h depuration. Asterisks above bars indicate statistically significant responses (*p<0.05, **p<0.01, ***p<0.001), while brackets indicate significant differences in PR between concentrations. Statistical analyses were performed using the Kruskal-Wallis test followed by Dunn’s *post hoc* test with Benjamin-Hochberg correction (α = 0.05).

In the absence of light stimulation (light OFF), swimming activity was markedly reduced compared to light conditions across all treatments, including controls, demonstrating a clear behavioral response to light stimulation. However, no significant difference was observed between GO4-exposed groups and controls under dark conditions. After 48 h of exposure, significant effects on mobility under light stimulation were observed, with swimming activity reduced by 27%, 51%, and 57% at GO4 concentrations of 0.7, 7, and 78 mg/l GO4, respectively, compared to controls. All treatments showed reduced mobility under dark conditions compared to light conditions, but no significant difference was observed between vehicle- and GO-treated groups in the absence of light stimulation, suggesting that GO compromises energy-demanding locomotor responses rather than basal mobility. These findings are in line with previous chronic exposure studies reporting that cardiac and locomotor impairments persist under prolonged GO exposure, with heart rate reductions of 19.4-26.9% observed after 24 days at concentrations as low as 0.625 mg/l [35]. Following depuration periods, *D. magna* showed partial recovery of swimming activity in response to light stimulation. After 24 h of exposure and 24 h of depuration, organisms exposed to 7 mg/l recovered to levels comparable to controls, whereas the 78 mg/l treatment group showed partial recovery but remained significantly impaired. After 48 h of depuration, swimming activity was partially restored at all concentrations under light stimulation, though organisms previously exposed to 78 mg/l still exhibited reduced mobility. In dark conditions, mobility remained consistently low across all treatments, including controls, throughout both depuration periods. The incomplete recovery at higher concentrations indicates that GO-induced locomotor and cardiac disruptions persist beyond the exposure period, consistent with observations that physiological endpoints, such as feeding activity, also show only partial restoration at elevated GO concentrations [36]. Such persistent impairments in cardiac performance and swimming capacity may increase vulnerability to predation and reduce foraging efficiency, potentially impacting population dynamics and food web structure in aquatic ecosystems [34–36].

Overall, the experiments carried out on the two species, *C. elegans* and *D. magna*, have clearly indicated that GO4 exhibits the greatest toxicity, as evidenced by both acute toxicity studies and behavioral deficits and sublethal alterations.

### Effect of GOs on plants

To systematically evaluate the phytotoxic effects of GO materials, the monocotyledonous rice (*Oryza sativa* L. Ky131) and the dicotyledonous pepper (*Capsicum annuum* L. Chao tianjiao) have been selected as model species, with experiments conducted via hydroponic systems and soil cultivation, respectively. Phytotoxicity was assessed using multi-dimensional indicators: germination rate and root length served as core phenotypic markers during seed germination, directly reflecting GO-induced inhibition of early plant development. Reactive oxygen species (ROS) and Malondialdehyde (MDA) levels were determined as oxidative stress biomarkers, with the former quantifying GO-induced free radical accumulation and the latter measuring cell membrane damage via lipid peroxidation. Chlorophyll and protein contents were used to characterize photosynthetic efficiency and cellular functional status, respectively, for evaluating GO’s impact on plant physiological functions. Antioxidant enzyme activities such as CAT, SOD, and peroxidase (POD) reflected plants’ defensive responses to oxidative damage, with their dynamic changes revealing potential regulatory pathways for GO toxicity. As illustrated in **Figures 8** and **9**, the potential phytotoxicity of GO was evaluated in a dose-dependent manner. During seed germination, GO materials exhibited pronounced concentration-dependent and material-specific effects on both rice and pepper. As shown in **Fig. 8A**, rice germination rates decreased to 6.7% under 0.1wt% GO1 and GO4 treatment, whereas GO2 and GO3 maintained germination rates at 46.7% and 40% even under the 0.4 wt% concentration, indicating no significant inhibitory effects. Pepper seeds showed lower sensitivity to GO, with germination rates maintained above 65% at 0.2 wt% concentration of GO1 (**Fig. 9A**), a concentration at which rice was greatly affected, highlighting the greater vulnerability of monocot early development to GO exposure. Furthermore, we did not observe a significant difference in toxicity between GO1 and GO4 in rice; however, it is evident that peppers exhibit a heightened sensitivity to GO4.

**Figure 8.**
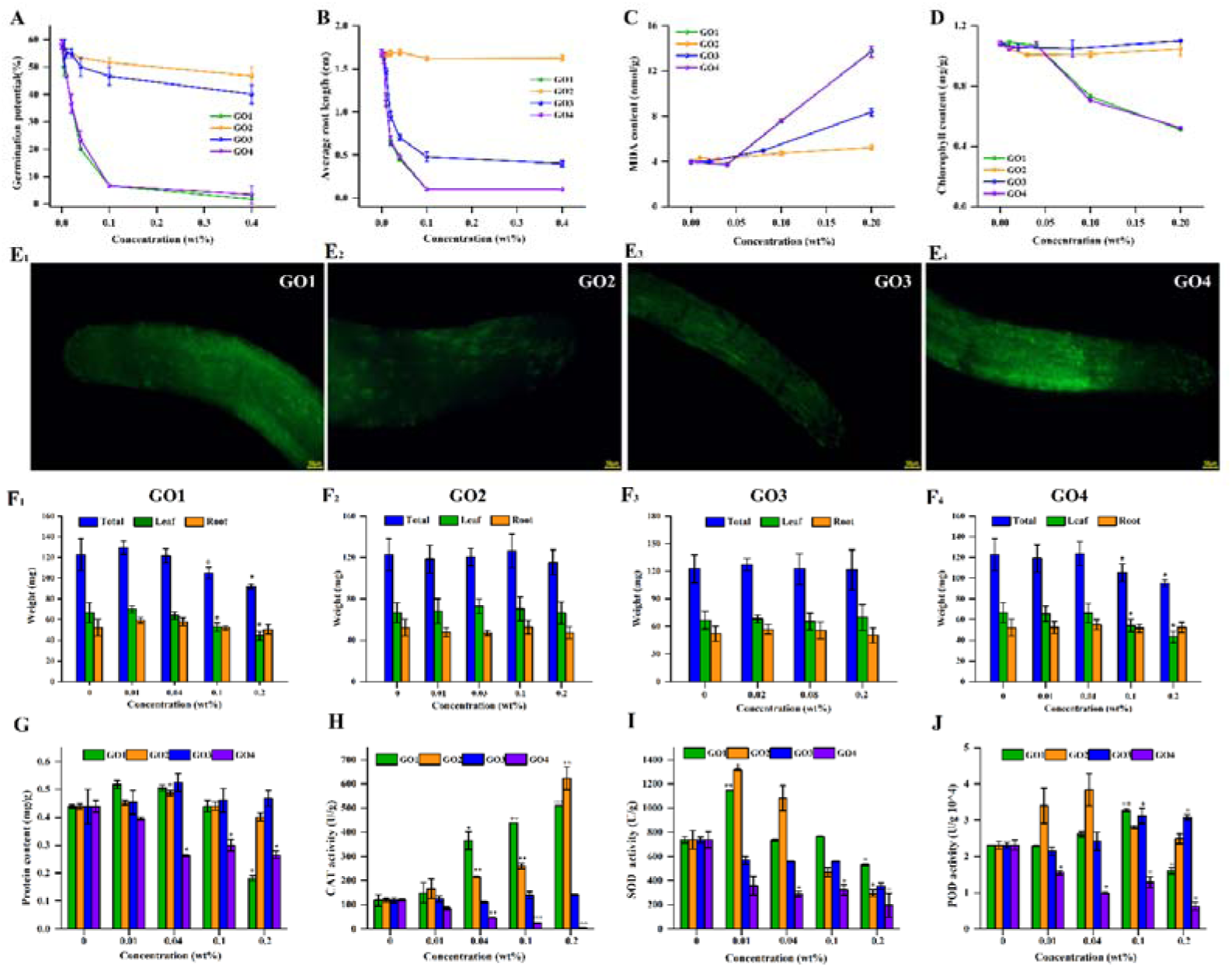
Effect of GO on the germination, physiological indices, and enzyme activity in rice. (A-B) Germination rate and root length of rice seeds exposed to GO1–GO4 at concentrations of 0, 0.01, 0.04, 0.1, and 0.2 wt%. (C) MDA content in rice roots. (D) Chlorophyll content in seedlings. (E) ROS fluorescence intensity in rice root tips exposed to 0.2 wt% GO1-GO4. (F) Total fresh weight, stem weight, and root weight. (G) Protein content. (H) Catalase, CAT; (I) Superoxide dismutase, SOD; (J) Peroxidase, POD in rice exposed to GO1-GO4. Data are expressed as mean ± SE. *p < 0.05, **p < 0.01 according to one-way ANOVA followed by LSD and Duncan’s *post hoc* tests.

**Figure 9.**
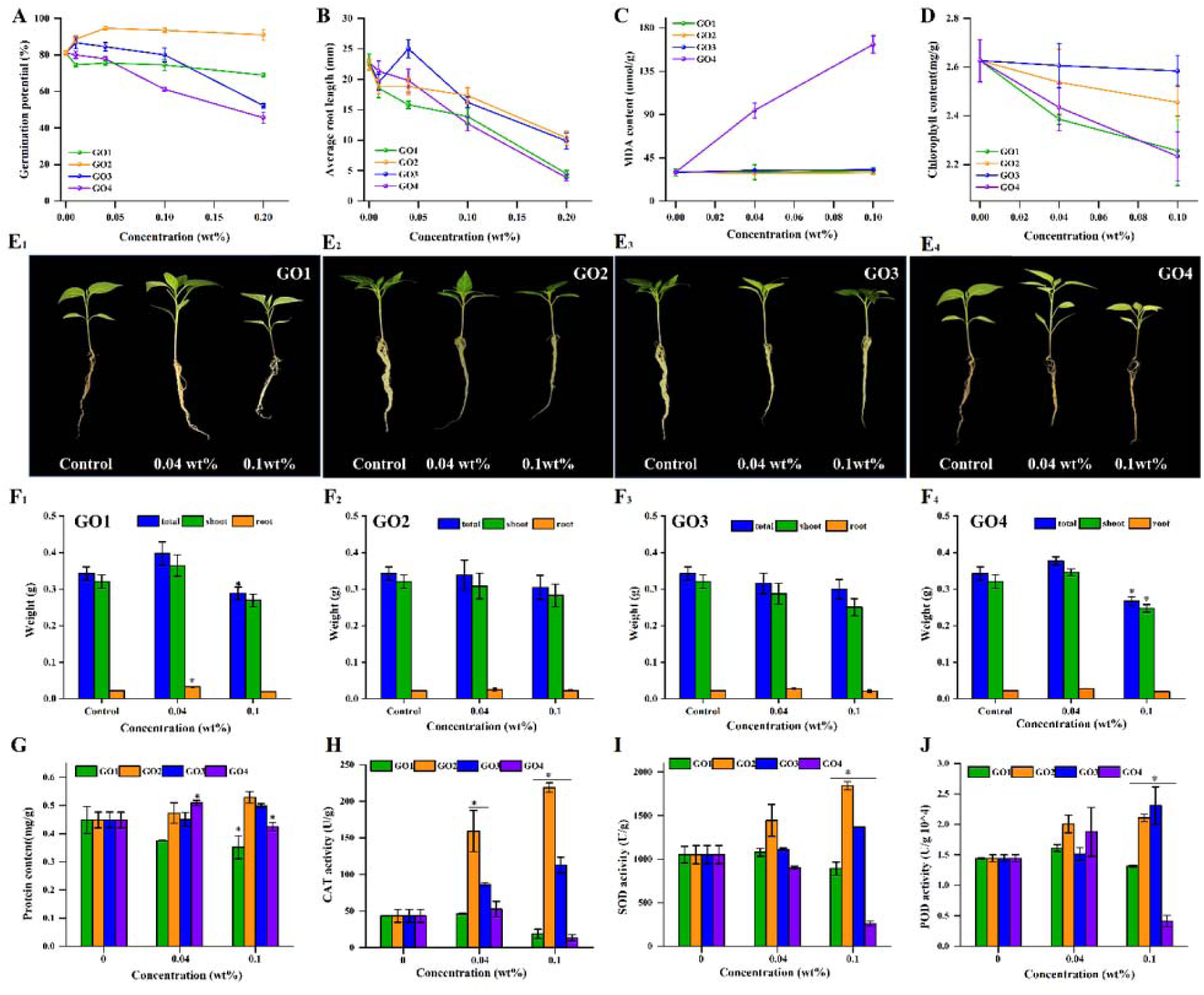
Effect of GO on the germination, physiological indices, and enzyme activity in pepper. (A-B) Germination rate and root length of seeds exposed to GO1–GO4 at concentrations of 0, 0.01, 0.04, 0.1, and 0.2 wt%. (C) MDA content in roots. (D) Chlorophyll content in seedlings. (E) Plants were grown in soil for 14 days by adding 0.04 wt% and 0.1 wt% nutrient solution. (F) Total fresh weight, stem weight, and root weight. (G) Protein content. (H) Catalase, CAT; (I) Superoxide dismutase, SOD; (J) Peroxidase, POD after exposure to GO1-GO4. Data are expressed as mean ± SE. *p < 0.05, **p < 0.01 according to one-way ANOVA followed by LSD and Duncan’s *post hoc* tests.

The dose-dependent inhibition of radicle growth was more pronounced: except for GO2, all other GOs caused progressive shortening of rice radicles with increasing concentrations, with GO1 and GO4 reducing root length to 0.1 cm at 0.1 wt% concentration (**Fig. 8B**). In pepper, radicle length at 0.04 wt% GO4 only showed a slight decrease, with exacerbation at higher concentrations (**Fig. 9B**). These phenotypic changes correlated closely with oxidative stress levels: ROS fluorescence intensity in rice root tips increased 2.1- and 2.7-fold under GO1 and GO4 (0.2 wt%) compared to the control (**Fig.8E**), while MDA content followed the toxicity order: GO4 ≈ GO1 > GO3 > GO2 (**Fig. 8C**), indicating a pattern of severity overlapping that found in animals. GO4 toxicity was also found in pepper, whereas MDA content in other GOs showed no significant change compared to the basal condition (**Fig.9C**). This indicates that high concentrations of GO4 disrupt cell membrane integrity via radical accumulation, thereby inhibiting radicle elongation. Notably, GO2 induced only a 14% increase in ROS levels, with no significant change in MDA at 0.2 wt%, suggesting that its low toxicity may be related to fewer surface C-O groups (hydroxyls, epoxides, and ethers).Physiological indices at the seedling stage further revealed complex impacts of GO materials on photosynthetic systems, protein synthesis, and antioxidant defense. In rice, GO1 and GO4 at 0.1 wt% reduced chlorophyll content by 32% and 35% (**Fig. 8D**), total fresh weight by 15% and 14%, and shoot weight by 18% and 19% (**Fig. 8F**), respectively. A similar trend was observed in pepper, with chlorophyll content decreasing by 15% under 0.1 wt% GO4 (**Fig. 9D**) and biomass changes mirroring those in rice (**Fig. 9F**).

Moreover, rice root protein content decreased significantly under 0.2 wt% GO1 and GO4 (**Fig. 8G**), whereas pepper exhibited a biphasic effect on protein content, with a 14% increase at 0.04 wt% followed by a 17% decrease at 0.1 wt% GO4 (**Fig. 9G**), aligning with phenotypic observations (**Fig. 9E**). Antioxidant enzyme activity assays revealed distinct modulations of oxidative stress markers by GO materials (**Fig. 8H-J**). Specifically, rice CAT activity increased at 0.2 wt% GO2 but decreased at 0.4 wt% GO2 (Fig. 8H), attributable to changes in surface epoxy group content, which modulate oxidative stress intensity. Rice SOD and POD displayed biphasic profiles, respectively peaking at 1.5× and 1.7× control levels at 0.04 wt% GO2 before declining by 60% at 0.2 wt% (**Fig. 8I-J**), reflecting initial antioxidant upregulation followed by high-concentration oxidative inactivation. In pepper, GO4 (0.1 wt%) induced over 50% suppression of CAT, SOD, and POD (**Fig. 9H-J**), whereas GO2/GO3 showed no significant decreases or only marginal increases, corroborating that GO4 is more toxic than the other forms. Collective analysis of multidimensional data establishes a toxicity paradigm for GO materials: oxidative stress, damage to membranes and proteins, and physiological dysfunction. At the cellular level, highly oxidized GO (notably GO1 and GO4) induces ROS overproduction in root tip cells via physical adsorption (**Fig. 8E**). Excessive free radicals initiate membrane lipid peroxidation, as evidenced by elevated MDA levels (**Fig. 8C**), thereby compromising membrane integrity, disrupting nutrient uptake, and impairing signal transduction. These perturbations directly manifest as suppressed seed germination and radicle elongation, with dose-dependent effects evident in both rice and pepper (**Figs. 8A and 9A**). At the physiological level, oxidative damage propagates to photosynthetic machinery, causing chlorophyll degradation (**Figs. 8D and 9D**) and subsequent reductions in biomass accumulation (**Figs. 8F and 9F**). The biphasic protein synthesis response (**Figs. 8G-J and 9G-J**) reflects a continuum of cellular adaptation: mild stress triggers compensatory protein upregulation, whereas severe stress induces a collapse of the synthetic pathway.

Through integrative analysis across developmental stages and plant taxa, these results delineate the “structure-activity” relationship of GO phytotoxicity, confirming oxidative stress as the conserved core toxic pathway spanning from seed germination to seedling development. GO4, characterized by a high density of carbonyl and epoxy functional groups, exhibits enhanced redox reactivity that drives more pronounced ROS accumulation at equivalent concentrations compared to other GO variants. Concurrently, the divergent responses between monocot rice and dicot pepper underscore the greater susceptibility of monocot early development to GO exposure, highlighting species-specific sensitivity as a critical modifier of nanomaterial toxicity.

Similar results have been reported in previous studies [37–40]. For instance, the germination rate of rice was significantly reduced at 100 mg/l, with a more pronounced inhibition observed at 500 mg/l [39, 41]. GOs at 100 and 250 mg/l significantly reduced rice biomass and inhibited shoot elongation, while MDA content was markedly increased. In contrast, no significant effects were observed for reduced GO [40].

Similarly, Xiao et al. [42] reported that the phytotoxicity toward *Brassica napus* L. followed the order GO > reduced GO > G-NH□, with these differences attributed to variations in surface chemical properties. In this study, GO4, bearing the thinnest particles, had the highest content of oxygen-containing functional groups, with significantly higher proportions of carbonyl and epoxy groups than those in partially reduced GO3 and highly reduced GO2. These carbonyl and epoxy groups may catalyze ROS generation via electron transfer reactions, which are considered the primary determinants of GO surface reactivity and provide active sites for interactions with biological interfaces [40, 43]. Additionally, the toxicity of various GOs varies with their aggregation behavior, size, and thickness in the aqueous phase [39, 44–47]. Greater dispersibility of GO is linked to higher phytotoxicity due to numerous oxygen-containing groups, such as carboxyl, hydroxyl, and epoxy, on its surface, which enhance both dispersibility and chemical reactivity. The tendency of GO2 to form large aggregates can restrict its movement and bioavailability in plants. Conversely, the higher dispersibility and surface charge density of GO4, its larger specific surface area, and greater reactivity can facilitate its cellular uptake and internal transport in plant tissues, ultimately resulting in increased phytotoxicity [48].

## Experimental

### GO synthesis

Four different types of GOs were synthesized using standardized protocols to obtain nanomaterials with different oxidation states and flake sizes (**Table 1**). GO1 (GO slurry) was synthesized using the patented method of Graphenea (San Sebastián, Spain, Patent EP3070053B1). Briefly, a modified Hummers method was employed to strongly oxidize graphite using KMnO4 and H2SO4. After purification, a 2 wt% slurry was obtained. GO2 (reduced GO) was synthesized using a method patented by Graphenea (US11981570B2). Briefly, a 4 mg/ml dispersion of GO1 was basified, and ascorbic acid was added as a reductant. After heating up and purifying the material, it was dried by atomization to obtain GO2 in powder format. GO3 (less oxidized GO) was obtained by changing the oxidant proportion of the reaction; the protocol was the same as the one used for the synthesis of GO1, but the graphite: KMnO4 ratio changed to 1:2 (w:w). After purification, a 2 wt% slurry of less-oxidized GO (GO3) was obtained. GO4 (smaller flake-size dispersion) was obtained by diluting GO1 to 0.4 wt% (4 mg/ml). After 5 min of stirring, the dispersion was subjected to 1 h of ultrasound treatment in an Elmasonic S80 system (Ruiselede, Belgium).

**Table 1.** List of the GO flake types, with the manufacturer’s characteristics of oxygen weight percentage (O%), flake size, and format.

| **GO** | **Format** | **%O** | **Flake size D<sub>50</sub> (µm)** | **Method** |
|---|---|---|---|---|
| GO1 | Water Slurry 20 mg/ml | 48.2 | 24.6 | Laser Diffraction |
| GO2 | Powder | 17.6 | 4.7 | Laser Diffraction |
| GO3 | Water Slurry 20 mg/ml | 35.9 | 18.9 | Laser Diffraction |
| GO4 | Water dispersion 4 mg/ml | 48.2 | 1.8 | Scanning electron microscopy |

### GO characterization

The physicochemical properties of the synthesized GOs were determined using various methods. SEM was conducted on a JEOL SEM JSM-IT800HL (Tokyo, Japan) equipped with an Energy Dispersive X-ray spectroscopy (EDS) system, ULTIM EXTREME SEM, with AZTEC software from OXFORD INSTRUMENTS PLC & SUBSIDIARIES (Abingdon, UK). Sample preparation for SEM acquisition is detailed in the SI.

The TEM experiments were conducted on a JEOL JEM-2100F UHR electron microscope (200 kV) equipped with an EDS system (Oxford UltimMax) and two Scanning Transmission Electron Microscopy (STEM) detectors: BF and High-Angle Annular Dark-Field (HAADF).

For AFM measurements, diluted samples were prepared according to SI and were analysed using a MultiMode AFM (Bruker, Milan, Italy). Data were acquired using the largest window (100 µm) scanner – J stage. AFM probes TESPA-V2 (Bruker) were used for all AFM measurements. NanoScope Analysis 2.0 (Bruker) was used to analyse AFM data.

XPS were recorded using the PHI XPS VersaProbe III energy spectrometer equipped with a monochromatic 1486.6 eV Al-Kα radiation source. A focused X-ray source with a beam size of 100 μm, a power of 25 W, and an e-beam energy of 15 kV was used. Charge neutralization was achieved using a complementary dual-beam method. The C 1s peak at 284.8 eV was used as a reference to calibrate acquired spectra. All spectra were processed using XPSPEAK41.

Raman spectra were acquired using an inVia Raman Microscopy system (Renishaw GmbH, Pleizhausen, Germany). A 532 nm laser line (1.29 mW power on stage) was used with a 50× L objective lens, 1800 lines mm-1 grating, and an extended 1024 charge-coupled device (CCD) array detector. Integration time was 10 s (5% of the maximum power), and each spectrum is the linear average of five distinct spectra collected from remote regions of each sample. Spectra were analysed using OriginLab. All bands were simulated using Lorentzian functions apart from the D” band, which was simulated using a Gaussian function following the method outlined in [13].

Molecular imaging was performed using a confocal Raman Alpha300 R microscope (WITec GmbH, Ulm, Germany) with a 532 nm laser, 50x and 100x magnification objectives (Zeiss, Germany), a 50 µm multimode fiber, and a CCD detector. Raman images (RI) were obtained pixel-wise in the x-y lateral plane using laser powers up to 30 mW and a 600 g mm-1 grating. The obtained spectra were further analysed using Project FIVE PLUS software (WITec GmbH), in which cosmic-ray correction, background subtraction, decomposition, and fitting of the recorded spectra were performed. All samples were filtered through Anodisc filters with a 20 nm pore size. The analysis was performed by combining high-quality information from RIs and BF-stitched images using a particle-searching algorithm in WITec Particle Scout 5.1. Raman spectra were recorded at every position of optically identified particles using autofocus alignment and were subsequently verified against data from the ST Japan Raman database. Each visualized particle (down to 260 nm in size) was analysed to determine its chemical composition, structural properties (e.g., D/G ratio), area, size, shape, and morphology. Thermogravimetric analysis (TGA) measurements were conducted on a TA Instruments TGA Discovery (New Castle, DE, USA). Samples were loaded into Pt pans and heated from 30 °C to 800 °C at 5 °C min-1 under a 20 ml/min N2 purge.

DCS was used to determine the dispersion of GOs. GO dispersions were prepared from dilutions of stock solutions (to 1 mg/ml) in the selected media. Prior to analysis, dispersions were homogenized by vortexing for 30 seconds and then injected into the DCS (except for stability-over-time evaluations). Particle density values for GO1/GO4, GO2, and GO3 were estimated as 1.36 g/ml, 1.80 g/ml, and 1.55 g/ml, respectively. The particle refractive index was 1.63, and the absorption coefficient (K) was 1. The non-sphericity factor was set to 3. A 8–24% sucrose gradient was employed, and polyvinyl chloride particles with a diameter of 544 nm served as standards. The centrifugation speed for the analysis was set at 16,000 rpm. Measurements were performed in triplicate, and the average data were reported as logWeight on apparent diameter. Data were reported as the maximum peak position ± half-width at half maximum (HWHM) to describe peak width, and ± SD to indicate data reproducibility. Only peaks with relative intensities greater than 10% of the maximum were considered noteworthy and reported. Comparing the peak positions obtained with the different dispersions allowed us to determine whether there were significant shifts in the particle-size distribution. Analysis of HWHM allowed us to assess whether peaks broaden, indicating a broader size distribution, whereas SD confirmed reproducibility.

To mimic the exposure of materials to biological fluids used *in C. elegans* experiments, GOs were dispersed in water collected from the C. elegans culture, which contained or did not contain OP50 *E. coli* (*Caenorhabditis elegans* Genetic Centre, CGC, University of Minnesota, Minneapolis, MN, USA) as nematode food. Samples were dispersed in ddH_2_O as a negative control. Samples were incubated at 20 °C for 24 hours and then centrifuged at 16,000 × *g* for 3 minutes to remove bacteria. Measurements were then performed as described above. To evaluate dispersion stability over time, a series of repetitive injections was performed at DCS. A total volume of 1 ml of dispersion was prepared in a 1.5 ml Eppendorf tube, vortexed for 30 seconds (t0), and then aliquots of 100 µl were sampled at different time points. The dispersions were left undisturbed between samplings. A reference point for sampling was established at 0.5 ml in the sample container to ensure consistent sampling. Results were plotted as log absorption versus apparent diameter and normalized to the intensity of the maximum peak at time 0, corresponding to the freshly prepared dispersion.

#### Caenorhabditis elegans

Bristol N2 *Caenorhabditis elegans* were obtained from the CGC and propagated at 20 °C on solid NGM seeded with OP50 *E. coli* for food. Age-synchronized animals were prepared by egg-laying. To this end, 30 gravid worms were transferred by picking on NGM plates seeded with OP50 *E. coli*. After 5 hours, the adult worms were removed, leaving only the eggs on the plates. The plates were incubated at 20 °C until the worms reached the L4 larval stage. To investigate the in vivo acute toxicity of GO, *C. elegans* were collected, centrifuged, and washed twice with ddH_2_O to remove bacteria (**Fig. S1**). Worms were then incubated for 2 hours at room temperature in orbital shaking, in the absence of *E. coli*, with increasing concentration (0.06-1 mg/ml) of GO (50 worms/50 µl). Worms were also incubated under the same experimental conditions with 50 μl of ddH2O as the negative control (vehicle) and 50 μl of 10 mM hydrogen peroxide as the positive control. After 2 hours of orbital shaking, worms were plated onto NGM plates seeded with OP50 *E. coli* and grown at 20 °C. Survival analysis was conducted from 24 to 72 hours after the GO treatment by counting the number of dead and alive worms on the plate. For each sample tested in the experiments, the number of viable worms, the number of dead worms, and the total number of worms were scored.

The effects of GOs on pharyngeal function and locomotor activity were assessed in synchronized worms at the L3 larval stage, fed 0.125 and 1 mg/ml GO as described above, at 2 and 24 hours post-treatment. The pharyngeal pumping rate was measured by counting the number of times the terminal bulb of the pharynx contracted in 1 minute (pumps/min). To measure locomotor activity, worms were picked up and transferred into a well of a 96-well ELISA plate containing 100 μl of 10 mM phosphate-buffered saline (PBS), pH 7.4, and their motility was evaluated by counting left–right movements per minute (body bends/min).

To determine the effect of GO on the lifespan and health span, synchronized *C. elegans* at the L3 larval stage were fed with GO at a concentration of 0.125 mg/ml, as previously described. Two hours after GO administration, 50 worms per condition were transferred to NGM agar plates seeded with OP50 *E. coli*; this was considered day 0 of the experiment. To prevent overlapping generations, nematodes were transferred daily to fresh NGM plates throughout the experiments. Survival was monitored daily, and worms were scored as alive, dead, or censored based on the following criteria: worms were considered alive if they displayed spontaneous movement or responded to gentle touch, and pharyngeal pumping was detectable; dead if they exhibited no movement or response to stimuli, and no visible pharyngeal pumping; and censored if they presented with an exploded vulva or were desiccated on the plate wall. Health span was evaluated in the same group of worms used for lifespan analysis by monitoring the progressive decline in locomotion. Worms were classified as moving if they displayed spontaneous locomotion; as not moving if they showed no movement or response to tactile stimuli; and as censored if they exhibited an exploded vulva or were found desiccated on the wall of the plate. For each sample, the number of viable, dead, moving, and non-moving worms, as well as censored worms, was recorded daily in a table. Kaplan–Meier statistical analysis was performed using the online software OASIS 2, and survival curves were generated.

The impact of GO on reproduction and larval development was assessed in N2-synchronized L4-stage worms fed GO at 0.125 mg/ml, as previously described. Two hours after treatment, worms were singly plated onto NGM agar plates seeded with OP50 *E. coli*, and the number of eggs laid by each worm was counted daily until the worm ceased laying. The larval development of eggs laid under these experimental conditions was monitored daily until the adult stage by counting the number of worms at each larval stage, as described in the literature [14]. Mitochondrial ROS production was assessed in live worms using MitoSOX™ Red staining (Thermo Fisher Scientific, Milan, Italy). Synchronized worms at the L3 larval stage were treated with GO at a final concentration of 0.125 mg/ml, as previously described. Negative control worms were treated with dH O, while positive controls were exposed to 1 mM H O. At the end of the treatment, 50 worms per condition were transferred onto freshly prepared 6 cm NGM plates containing 10 μM MitoSOX™ Red and seeded with UV-killed OP50 *E. coli*. After incubation for 16 h in the dark at 20 °C, worms were transferred to new NGM plates seeded with live OP50 *E. coli* for 1 h to remove residual dye from the pharyngeal lumen and gut. For imaging analysis, worms were mounted on glass slides coated with 2% agarose, anesthetized with 10 mM levamisole (Sigma-Aldrich Co., Milan, Italy), and fixed with ProLong™ Glass Antifade Mountant (Thermo Fisher Scientific). Slides were covered with coverslips, and fluorescence signals were analysed using an inverted fluorescence microscope Olympus U-RFL-T (Olympus Italia S.r.l., Segrate, Italy). Virtual slide images were acquired with the Olympus VS120 virtual slide microscope. For each experimental condition, 10 worms were analysed, and images of the pharynx and body wall muscles were acquired. The integrated fluorescence intensity (integrated density) was quantified using Fiji software.

#### Daphnia magna

*Daphnia magna* (Straus), originating from a clonal lineage, was used. Parthenogenetic cultures were established in 1 l aquaria, each containing 40 individuals, and maintained at 20 °C in a medium composed of 100% Volvic® water supplemented with thiamine, cyanocobalamin, and biotin, under a 16:8 h light: dark photoperiod. The medium was renewed three times per week. Daphnids were fed daily with a mixed algal diet composed of Raphidocelis subcapitata (5 × 10□cells/*Daphnia*/day), Desmodesmus subspicatus (2 × 10□cells/*Daphnia*/day), and Chlorella vulgaris (2 × 10□cells/*Daphnia*/day), corresponding to an estimated carbon input of 0.09 mg C/daphnid/day. To ensure the physiological integrity and sensitivity of the cultures, routine quality-control tests using K□Cr□O□were performed monthly in accordance with ISO 6341 [15] and OECD Test Guideline 202 (OECD, 2012) [16]. All test organisms used in the experiments met the required sensitivity range for the reference toxicant, with an EC_50_ of 0.9 mg/l (accepted EC_50_ = 0.6 – 2.1 mg/l), confirming their suitability for toxicological assays.

Acute immobilization assays (48 h) were performed according to OECD guideline 202 (OECD, 2012) [16] and ISO 6341 (ISO, 2012) [15] at 20 ± 2 °C in full darkness. Neonates (<24 h old) from 2–6-week-old parent *D. magna* were used without feeding throughout the exposure period. The test medium (ISO formulation) consisted of 294 mg/L CaCl□x 2 H□O, 123 mg/l MgSO□·7 H□O, 64.7 mg/l NaHCO□, and 5.75 mg/l KCl. Preliminary range-finding assays using seven GO concentrations (7.8–1000 mg/l) were conducted to establish suitable test concentrations. Initial experiments compared pH 8 (standard ISO/OECD conditions) with pH 6 to reduce pH drift during exposure while ensuring organism survival. For definitive testing, daphnids were exposed to 1–500 mg/l GO and to one control: test medium alone (**Fig. S2**). Two independent experiments were conducted for each GO material. Organism immobilization was recorded at 24 and 48 h. The pH was monitored in control vessels at 0 and 48 h and in selected treatment concentrations at 48 h.

For the sub-acute test, *D. magna* neonates (< 24 h old) were exposed to 0, 0.7, 7, and 78 mg/l GO4, with 78 mg/l corresponding to the EC50 value previously determined in the acute toxicity test. Exposures were conducted for 24 and 48 h. Following 24 h of exposure, subsets of individuals were transferred to clean medium for depuration periods of 24 h or 48 h (**Fig. S2**). All exposures and depuration phases were conducted in complete darkness and in the absence of food. Each condition was tested in 5 biological replicates, with five individuals per replicate (N = 25 per condition). Survival, heartbeat rate, and behavioral activity were assessed at each time point, both immediately after exposure and following removal of the solution containing GO4. At each time point, survival rates were calculated.

The heartbeat rate was also determined. At each time point, 10 individual *D. magna* were placed in separate wells of a 6-well plate containing ISO medium. For each, heartbeats were manually counted over a 30-second interval using a manual cell counter. Each measurement was repeated 3 times per individual to ensure consistency.

For each treatment, behavioral swimming activity was measured in a pool of 12 organisms. Assays were performed using a Daniovision system, including a Daniovision observation chamber (DVOC) with near-infrared illumination, coupled with a temperature-control unit that maintained the plates at 20 °C. The automated delivery of visible-light stimuli from the DVOC was controlled by EthoVision XT v17 software (Noldus, Wageningen, the Netherlands) [17]. Trials were conducted on a 24-well plate so that the movement of *D. magna* would not be restricted. One organism/well and 12 wells per condition were used. Before delivering the first light stimuli, organisms were left in the DVOC for 20 min in full darkness to acclimate. The test consisted of a white-light routine with 15 min of light at 50% intensity (290 lux), followed by 5 min of 100% darkness. The white-light routine was repeated 3 times at each time point [17, 18]. Videos were recorded at 25 FPS. Total distance (mm) and velocity (mm/s) were analyzed using the multitracking module of the Ethovision software [17, 18].

### Plants

To evaluate the effect of GOs on seed germination and root elongation, rice and chili pepper seeds were purchased from Jinyuliang Seed Co., Ltd., Vigorous (Shenyang, Liaoning, China). Uniformly sized rice and chili pepper seeds were selected, disinfested with sodium hypochlorite, rinsed thoroughly with clean water, and placed in culture media soaked with filter paper for germination. Each culture medium contained 20 seeds, each treated with a different GO concentration (0-0.2 wt%, corresponding to 0-2.0 mg/ml for GO1, GO3, and GO4, and 0-0.4 wt%, corresponding to 0-4.0 mg/ml for GO2). The treated Petri dishes were incubated in a constant-temperature, constant-humidity chamber, in the dark, for 7 days. The germination conditions of the seeds were recorded daily, and the incubator was opened once daily to replenish the culture solution with an appropriate amount of ultrapure water to compensate for evaporation and maintain a constant culture solution concentration. After 7 days, the germinated seeds were harvested to measure root elongation and calculate germination rate. Germinated seeds of uniform emergence were transferred to hydroponic/potting containers for cultivation, with 6 plants per container. Upon reaching the two-leaf stage, the seedlings were treated with varying concentrations of GOs. After 7-14 days of cultivation, their physiological parameters (phenotype, plant height, root length) were measured. After cleaning the seedlings and drying them on filter paper, the plants were separated into aboveground and belowground parts, and their fresh weights were measured individually. Each treatment was replicated three times. All experiments were conducted under conditions of daytime temperature of 28°C, nighttime temperature of 22°C, a 16-hour light period and a 12-hour dark period, relative humidity of approximately 70%, and a light intensity of 3000 Lux.

Plant root enzyme activities were measured after treatment with different concentrations of GOs using kits from Solarbio Biotechnology Company (Beijing, China). In particular, the activity of the following enzymes was evaluated: Catalase (CAT, cod. BC0205), superoxide dismutase (SOD, cod. BC5165), peroxidase (POD, cod. BC0095), and Malondialdehyde (MDA, cod. BC0025). Chlorophyll Assay Kit (cod. BC0995) was used to determine the chlorophyll content in plant leaves. Each treatment was replicated three times.

To assess oxidative damage, the 2’, 7’-dichlorofluorescein (H2DCF-DA) fluorescent probe (Coolaber, Beijing, China) was used to detect ROS activity. A 10 mg/ml H2DCF-DA stock solution was prepared in DMSO and further diluted to a working concentration of 10 µg/ml. Root tips from seedlings subjected to different treatments were first cleaned 2–3 times with 5 mM PBS, then incubated in the 10 µg/ml H2DCF-DA working solution for 1 hour in the dark. The root tips were then rinsed 2–3 times with 5 mM PBS to remove excess dye, and fluorescence was observed and imaged with a fluorescence microscope (Olympus CKX53, Tokyo, Japan) at 485 nm excitation.

### Statistical analysis

*C. elegans* data were analysed using GraphPad Prism 10.0 (San Diego, CA, USA). Data are presented as mean ± SEM. Median inhibitory concentration (IC_50_) was analysed by nonlinear regression analysis. Normality was assessed, and because the distribution was normal, statistical analysis was performed using one-way ANOVA followed by Bonferroni’s *post hoc* test. Statistical significance was set at p < 0.05.

Data obtained from *D. Magna* experiments were analyzed using R software (version 4.3.2, 2023) with RStudio (version 2023.12.1, build 402). Median effective concentrations (EC_50_) and their 95% confidence intervals for *D. magna* acute immobilization were calculated using concentration-response curve fitting with the drc package in R. Statistical differences between EC values were determined based on non-overlapping 95% confidence intervals. For the subacute test, all data are presented as mean ± SD. Normality was assessed using the Shapiro-Wilk test, and homogeneity of variance was evaluated using Levene’s test. Due to non-normality and/or heterogeneity of variances, a nonparametric analysis was performed using the Kruskal-Wallis test, followed by Dunn’s *post hoc* test for pairwise comparisons. Statistical significance was set at p < 0.05. Plant data were analysed using IBM SPSS Statistics, and figures were plotted using Origin 2024. Data are presented as mean ± SE. Statistical analysis was performed using one-way ANOVA followed by LSD and Duncan’s *post hoc* tests for multiple comparisons. Statistical significance was set at p < 0.05.

## Conclusions

This study clarifies the relationship between the physicochemical properties of industrial nanoflake GOs and their impact on animal and plant toxicity. The oxidation state resulted as a key determinant of toxicity. GO2, the most reduced sample with the lowest oxygen functional content, exhibited the least acute and subacute effects in both animals and plants. It retained the strongest π-π interactions, as inferred by the sizeable restoration of the graphitic percolation network.

Unlike the other GOs, the strong π-π interparticle interactions in GO2 overwhelm the strong yet scarce hydrogen bonding, causing particle aggregation regardless of the ionic strength of the stabilizing biological media. Only GO2 particles consistently exhibit a very defective and crumpled morphology while having the smallest lateral size and the largest thickness among all GOs. The hydrophobic and morphological characteristics of GO2 play critical roles in defining its stability in different media. GO2 particles are three-dimensional, rigid, very defective, and possess sharp peripheries. GO3 represents a transitional variant in the oxidation process, exhibiting a hybrid character between the highly oxidized GO1/GO4 and the graphitic reduced GO2. While GO1 and GO4 maintain high oxygen functional content and consequently wide interlayer spacings, GO3 exhibits a reduced interlayer spacing and a moderate C/O ratio. It is therefore intuitive to assume that GO3 dispersibility and fate in biological media manifest somewhere between GO1/GO4 and GO2. These findings, indicating increased aggregation of GO2 relative to GO1, GO3, and GO4, suggest that the lower toxicity of GO2 can be attributed to its limited capacity to cross biological membranes in both plant and animal models. In contrast, the higher stability of GO4, resulting from its smaller size and high oxygen functional content, could promote internalization and consequently enhance its toxicological impact. However, potential contributors to this toxicity include further oxidation of C–O groups to C=O groups, forming carbonyl/quinone species [49], as well as the possible leaching of residual sulfur species from the oxidation process used to obtain GO1.

In *C. elegans*, widely used for assessing the biological effects of nanomaterials and environmental pollutants, a single administration of the three oxidized materials, GO1, GO3, and GO4, but not the reduced GO2, caused acute and subacute toxic effects. GO4 was the most toxic material for the worms, with the lowest IC_50_ value and the greatest impact on the lifespan and health span of nematodes. In addition, it is the material with the greatest pro-oxidant activity, as indicated by its effect on mitochondrial ROS production. Notably, the toxic effects of GO1, GO3, and GO4 are permanent and affect development, suggesting that the dispersion of these materials in the environment could significantly delay the emergence of new nematodes. Interestingly, the worm’s development was also affected by GO2, suggesting that this effect is unrelated to the oxidative state and flake size and raising concerns about its safety for animal health.

In *D. magna*, GO4 was also the most toxic material, with the lowest EC_50_. The acute toxicity ranking: GO4 > GO1 ≥ GO3 > GO2 demonstrated differential effects among GO variants, consistent with observations in *C. elegans*. This cross-species consistency suggests that the physicochemical properties of GO, particularly oxidation state and flake size, influence toxicity across different invertebrate models. Sub-lethal endpoints revealed greater sensitivity than immobilization: reductions in heart rate and concentration-dependent decreases in swimming activity under light stimulation indicate that physiological and locomotor functions are early targets of GO toxicity.

Highly oxidized GO4 also exhibited the strongest plant toxicity, while the most reduced GO2 showed the lowest toxicity. Physio-biochemical analyses revealed that a burst of ROS was directly induced by highly oxidized GOs in root tip cells, inducing membrane lipid peroxidation. Consequently, seed germination and radicle elongation were inhibited, while plant growth inhibition, leaf photosynthetic damage, and biomass reduction were aggravated. Although a higher reactivity toward biomolecules would be expected for the more reduced GO2, we propose that its low toxicity is primarily associated with extensive aggregation and restricted translocation within organisms.

Overall, the results of this study underscore the intrinsically multidimensional nature of industrially produced GO used in diverse applications and the key role of physicochemical properties in its potential toxicity. The unprecedented platform applied herein could be employed to parameterize other nanomaterials and toxicity descriptors to assess their potential impact in the (eco)toxicological field and to establish a robust level of safety.

## Author contributions

A. Subrati: Data Curation, Formal Analysis, Investigation, Methodology, Writing – original draft, Writing – review & editing. P. Bigini: Conceptualization, Data curation, Resources, Funding acquisition, Writing – original draft, Writing – review & editing, Project administration. A. De Luigi: Data Curation, Formal Analysis, Investigation, Methodology. C. Natale: Data Curation, Formal Analysis, Investigation, Methodology, Writing-original draft. K. Mehennaoui: Data Curation, Formal Analysis, Investigation, Methodology, Writing-original draft. Marianne Houin: Data Curation, Formal Analysis, Investigation, Methodology. Aline Chary: Data Curation, Formal Analysis, Investigation, Methodology, Writing-original draft. T. Serchi: Conceptualization, Methodology, Funding acquisition, Writing – review & editing. X. Zhang: Data Curation, Formal Analysis, Investigation, Methodology, Writing-original draft. K. Zhao: Data Curation, Formal Analysis, Investigation, Methodology. B. Alonso; Data Curation, Investigation, Methodology, Resources, Writing-original draft. T. Venus: Data Curation, Formal Analysis, Investigation, Methodology, Writing-Original Draft. A. Saorin: Data Curation, Formal Analysis, Investigation, Methodology. M.P. Monopoli: Conceptualization, Data curation, Resources; I. Estrela-Lopis: Conceptualization, Methodology, Funding acquisition, Writing – original draft, Writing – review & editing. C. Gao: Conceptualization, Methodology, Funding acquisition, Writing – original draft, Writing – review & editing. A. Zabeo: Writing – original draft, Writing – review & editing; D. Hristozov: Writing – original draft, Writing – review & editing; S. Moja: Conceptualization, Data curation, Resources, Funding acquisition, Writing – original draft, Writing – review & editing, Project administration. L. Diomede: Conceptualization, Data curation, Resources, Funding acquisition, Writing – original draft, Writing – review & editing, Project administration.

## Data availability

Data for this article are available at Zenodo upon request to the corresponding authors.

## Acknowledgements

This research was funded by the European Union’s Horizon Europe RESEARCH AND INNOVATION ACTIONS PROGRAMME, Project “Platform Optimisation to Enable Nanomaterial Safety Assessment for Rapid commercialization (POTENTIAL)”, Grant number 101092901. CG reports that financial support was provided by the National Key Research and Development Program of China (2023YFE0108700). LD and CN acknowledge GCG, which is funded by the NIH Office of Research Infrastructure Programs (P40 OD010440), for providing Caenorhabditis elegans and OP50 Escherichia coli.

