## Supplementary data for "The key role of oxidation state and flake size in the ecotoxicological impact of industrial graphene oxide nanoflakes"

**Supplementary Method**

Scanning Electron Microscopy of Graphene Oxide

This method provides a stepwise procedure for obtaining particle-size and aspect-ratio descriptors. It covers sample preparation and analysis.

The protocol is described in detail on protocols.io,

[DOI: https://dx.doi.org/10.17504/protocols.io.ewov19opklr2/v1 (Private link for reviewers: https://www.protocols.io/private/A167F8092FD011EF83050A58A9FEAC02 to be removed before publication.) and is included for printing as SI File with this article.

**Supplementary Figures**

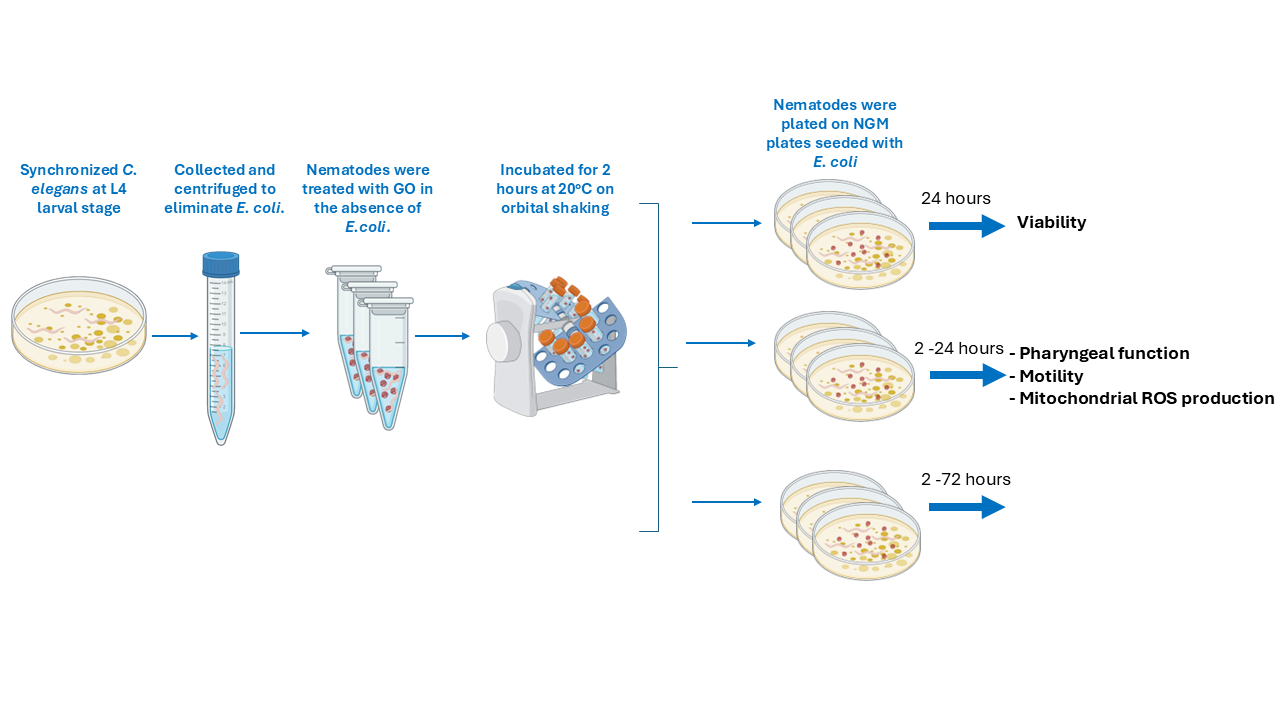
**Figure S1.** Schematic representation of the experimental design applied for investigating the impact of GOs on *C. elegans*.

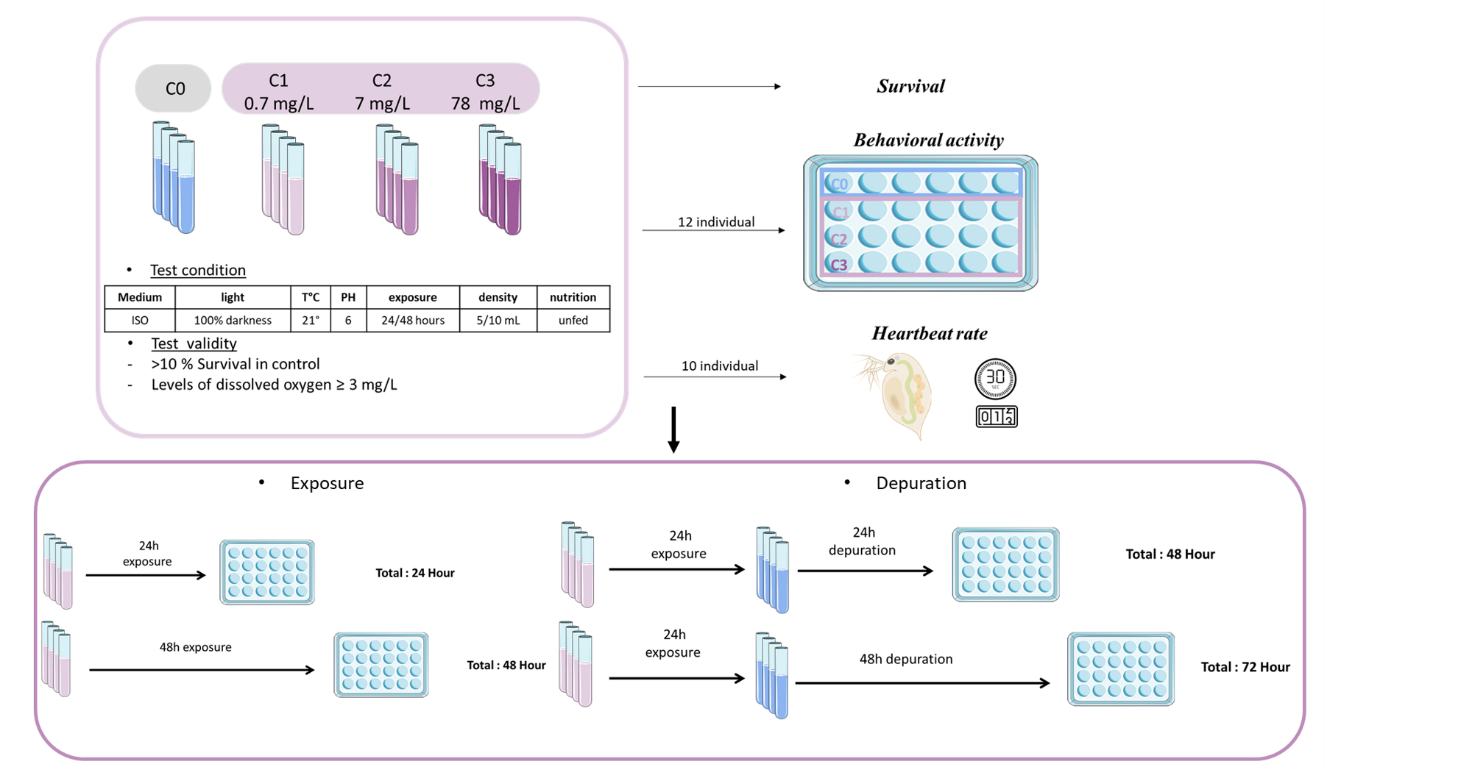

**Figure S2.** Schematic representation of the experimental design applied for investigating the impact of GOs on *Daphnia magna*.

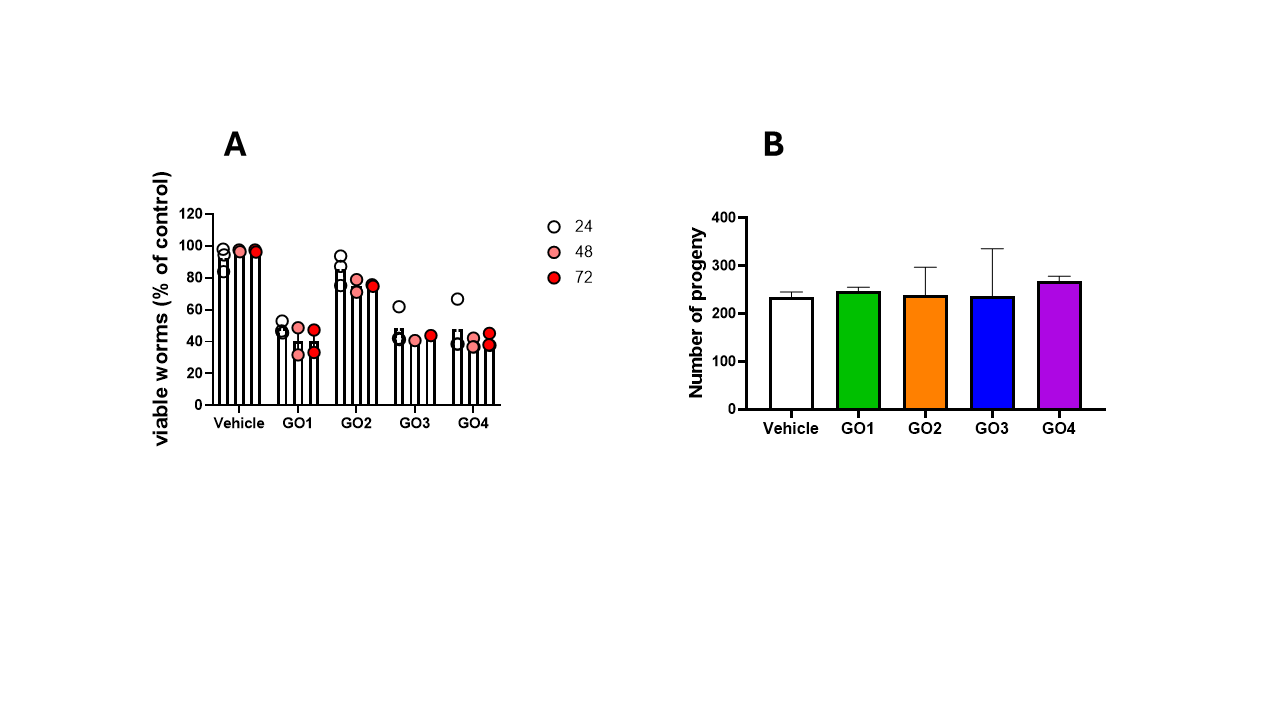

**Figure S3.** **Effect of GOs on *C. elegans* viability and reproduction. A**) Synchronized worms were fed with 1 mg/ml of different GOs suspended in water without OP50 *E. coli* (50 worms/50 µl). Worms were fed water alone (Vehicle) (50 worms/50 µl) as a negative control. After 2 h of incubation at 20°C, worms were plated on NGM plates seeded with fresh bacteria, and the number of viable worms was assessed at 24, 48, and 72 h. Data are the mean of viable worms (% of vehicle) ± SEM (N=50) from 3 independent experiments. **B**) Effect of GOs on the development of nematodes. Worms (50 worms/50 μl) were fed for 2 hours with 0.125 mg/ml GOs suspended in water, without *OP50 E. coli*, then plated on NGM plates seeded with the bacteria. Control worms were fed with only water (vehicle). The number of progenies after plating was scored until the worm stopped laying. Mean ± SEM (N = 12).

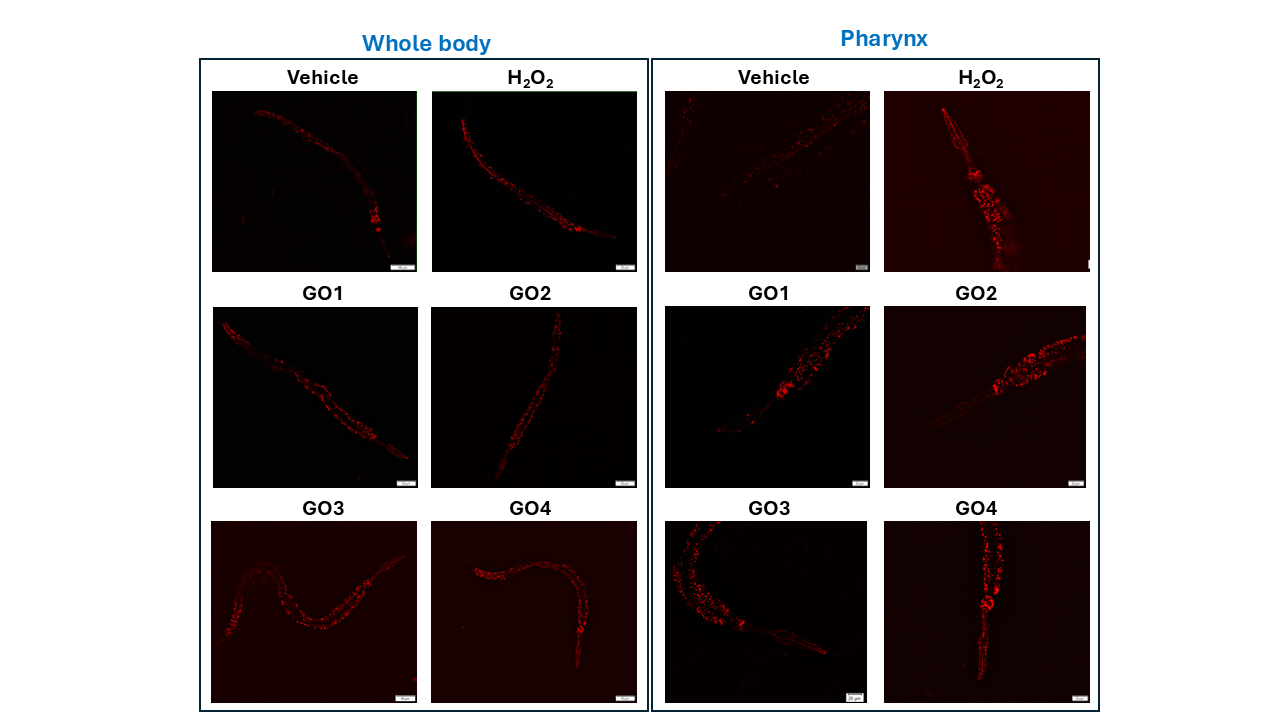

**Figure S4.** **Effect of GOs on mitochondrial superoxide production.** Representative images of mitochondrial superoxide production, detected with MitoSOX™ Red, in the whole body and the pharynx of worms treated with 0.125 mg/ml GOs for 24 h or with 1 mM H_2_O_2_ for 30 minutes, as a positive control. Scale bars: = 50 µm for whole body and 20 µm for pharynx.

**Table S1**. Summary of particle analysis by SEM. Numbers in brackets are of the area-weighted cumulative particle size distribution.

| **Sample** | **Size (µm)** | | | **Aspect ratio** | | | **SEM image** |
| --- | --- | --- | --- | --- | --- | --- | --- |
|  | **D10** | **D50** | **D90** | **D10** | **D50** | **D90** |  |
| GO1 | 1.2  (**7.7**) | 4.4  (**24.8**) | 16.7  (**66.5**) | 1.3 | 1.8 | 3.1 | 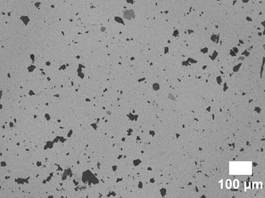 |
| GO2 | 1.1  (**4.8**) | 3.1  (**9.7**) | 10.0  (**18.1**) | 1.2 | 1.5 | 2.0 | 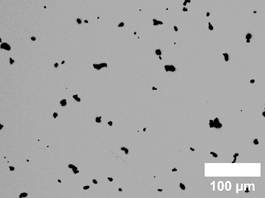 |
| GO3 | 2.0  (**6.8**) | 7.2  (**13.9**) | 15.2  (**33.2**) | 1.2 | 1.6 | 2.2 | 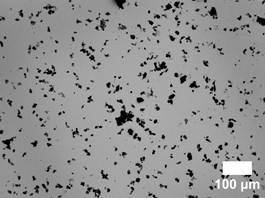 |
| GO4 | 1.5  (**5.6**) | 4.2  (**17.2**) | 14.8  (**35.2**) | 1.2 | 1.4 | 1.6 | 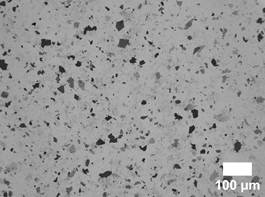 |

**Table S2**. Survey, wide scan XPS, and element quantification of samples GO1, GO2, GO3, and GO4. Survey- and high-resolution-obtained atomic carbon-to-oxygen ratio (C/O).

| **Sample** | **Content (at. %)** | | | | **C/O determined by** | |
| --- | --- | --- | --- | --- | --- | --- |
|  | **C** | **O** | **N** | **S** | **Survey** | **High-resolution** |
| GO1 | 70.20 | 27.70 | 0.70 | 1.40 | 2.53 | 2.81 |
| GO2 | 90.00 | 9.74 | 0.22 | 0.04 | 9.24 | 9.16 |
| GO3 | 75.08 | 23.02 | 0.39 | 1.51 | 3.26 | 3.58 |
| GO4 | 70.62 | 27.50 | 0.53 | 1.35 | 2.57 | 2.80 |

**Table S3**. High-resolution, narrow scan, C 1s X-ray photoelectron spectra deconvolution summary. BE is binding energy. At. % is the atomic contribution of the component.

| **Sample** |  | **C=C** | **C‒C** | **C‒O** | **C=O** | **COO** | **π-π*** |
| --- | --- | --- | --- | --- | --- | --- | --- |
| GO1 | BE (eV) | 284.70 | 285.24 | 286.80 | 288.16 | 289.07 | 290.90 |
|  | At. % | 30.03 | 13.62 | 43.93 | 5.84 | 4.07 | 2.50 |
| GO2 | BE (eV) | 284.79 | 285.40 | 286.74 | 287.90 | 289.00 | 291.04 |
|  | At. % | 46.00 | 25.83 | 10.12 | 2.07 | 5.27 | 10.71 |
| GO3 | BE (eV) | 284.78 | 285.31 | 286.81 | 287.90 | 289.00 | 290.90 |
|  | At. % | 36.79 | 16.55 | 31.70 | 5.07 | 5.26 | 4.63 |
| GO4 | BE (eV) | 284.71 | 285.20 | 286.80 | 288.16 | 289.14 | 290.96 |
|  | At. % | 30.78 | 11.83 | 45.25 | 7.00 | 3.33 | 1.81 |

**Table S4.** Peak integrated intensity (area) ratios calculated from the deconvolution results of the Raman spectra of the GO materials. L_a_ and n_D_ are the crystallite lateral width and defect density.

| **Sample** | **A_D_/A_G_** | **A_D'_/A_G_** | **A_D_/A_D'_** | **L_a_ (nm)** | **n_D_ ×10^11^ (defect/cm^2^)** |
| --- | --- | --- | --- | --- | --- |
| GO1 | 3.10 | 0.36 | 8.58 | 6.20 | 9.30 |
| GO2 | 2.37 | 0.23 | 10.22 | 8.12 | 7.09 |
| GO3 | 2.98 | 0.29 | 10.29 | 6.44 | 8.94 |
| GO4 | 3.14 | 0.39 | 7.94 | 6.13 | 9.39 |

**Table S5**. Deconvolution results of the Raman spectra of the GO materials.

| **Sample** |  | **Band** | | | | | | | | |
| --- | --- | --- | --- | --- | --- | --- | --- | --- | --- | --- |
|  |  | **D*** | **D** | **D"** | **G** | **D'** | **M** | **2D** | **D+D'** | **2D'** |
| **GO1** | Intensity | 86 | 2357 | 508 | 1827 | 1086 | 71 | 324 | 497 | 150 |
|  | FWHM (cm^-1^) | 73 | 78 | 134 | 32 | 20 | 30 | 170 | 155 | 75 |
|  | Area (cm^-1^) | 19602 | 573910 | 85577 | 184939 | 66918 | 6790 | 173288 | 242797 | 35252 |
|  | Center (cm^-1^) | 1116 | 1353 | 1530 | 1585 | 1608 | 1772 | 2682 | 2931 | 3176 |
| **GO2** | Intensity | 72 | 1586 | 251 | 1244 | 553 | 17 | 210 | 247 | 76 |
|  | FWHM (cm^-1^) | 113 | 53 | 138 | 29 | 15 | 29 | 142 | 98 | 67 |
|  | Area (cm^-1^) | 25562 | 263925 | 43585 | 111517 | 25812 | 1569 | 93318 | 75755 | 15878 |
|  | Center (cm^-1^) | 1176 | 1346 | 1518 | 1583 | 1611 | 1771 | 2682 | 2926 | 3164 |
| **GO3** | Intensity | 34 | 1521 | 346 | 1212 | 629 | 32 | 200 | 285 | 98 |
|  | FWHM (cm^-1^) | 48 | 64 | 127 | 27 | 15 | 57 | 130 | 112 | 93 |
|  | Area (cm^-1^) | 5214 | 303413 | 55082 | 101691 | 29476 | 5741 | 81707 | 100249 | 28551 |
|  | Center (cm^-1^) | 1121 | 1352 | 1536 | 1585 | 1610 | 1796 | 2691 | 2927 | 3168 |
| **GO4** | Intensity | 76 | 2027 | 454 | 1576 | 974 | 70 | 345 | 386 | 170 |
|  | FWHM (cm^-1^) | 111 | 78 | 139 | 32 | 20 | 42 | 195 | 116 | 99 |
|  | Area (cm^-1^) | 26582 | 495050 | 79077 | 157903 | 62316 | 9221 | 211265 | 140862 | 52698 |
|  | Center (cm^-1^) | 1116 | 1354 | 1530 | 1585 | 1610 | 1776 | 2706 | 2937 | 3166 |

**Table S6.** List of the main physicochemical features.

|  |  | **GO1** | **GO2** | **GO3** | **GO4** |
| --- | --- | --- | --- | --- | --- |
| **Format** |  | Slurry, 20 g/L | Fine powder | Slurry, 20 g/L | Dispersion, 4 g/L |
| **CHNS elemental analysis (wt.%)** | C | 48.7 | 82.0 | 60.9 | 48.7 |
|  | H | 0.6 | 0.2 | 0.2 | 0.6 |
|  | N | 0.1 | 0.2 | 0.1 | 0.1 |
|  | S | 2.4 | 0.0 | 2.9 | 2.4 |
|  | O | 48.2 | 17.6 | 35.9 | 48.2 |
| **XPS elemental analysis (at. %)^[a]^** | C | 70.2 | 90.0 | 75.1 | 70.6 |
|  | N | 0.7 | 0.2 | 0.4 | 0.5 |
|  | S | 1.4 | 0.0 | 1.5 | 1.4 |
|  | O | 27.7 | 9.7 | 23.0 | 27.5 |
| **C/O (at./at.)^[b]^** |  | 2.81 | 9.16 | 3.58 | 2.80 |
| **Defect density, A_D_/A_G_^[c]^** |  | 3.10 | 2.37 | 2.98 | 3.14 |
| **Defect type, A_D_/A_D'_^[c]^** |  | 8.58 | 10.22 | 10.29 | 7.94 |
| **Particle size (µm)^[d]^** | D10: | 7.7 | 4.8 | 6.8 | 5.6 |
|  | D50: | 24.8 | 9.7 | 13.9 | 17.2 |
|  | D90: | 66.5 | 18.1 | 33.2 | 35.2 |
| **Particle aspect ratio** | D10: | 1.3 | 1.2 | 1.2 | 1.2 |
|  | D50: | 1.8 | 1.5 | 1.6 | 1.4 |
|  | D90: | 3.1 | 2.0 | 2.2 | 1.6 |
| **Particle thickness (nm)** | D10: | 9.5 | 92.7 | 9.0 | 8.5 |
|  | D50: | 65.6 | 156.6 | 45.6 | 20.0 |
|  | D90: | 252.6 | 393.0 | 177.9 | 82.4 |
| **Fraction of particles < 1 µm (%)^[e]^** |  | 19 | 2 | 23 | 33 |
| **Interlayer spacing (Å)** |  | 8.0 | 3.5 | 6.5 | 8.1 |
| **Crystal c width, L_c_ (Å)** |  | 149.1 | 13.0 | 44.0 | 137.4 |
| **Crystal ab width, L_a_ (Å)** |  | 62.0 | 81.2 | 64.4 | 61.3 |
| **Oxygen-functionalized content (wt.%)^[f]^** |  | 53.4 | 18.8 | 41.8 | 53.4 |
| ^[a]^Elemental analysis determined by XPS survey spectra (wide scan).  ^[b]^Degree of oxidation descriptor determined from high-resolution XPS spectra (narrow scan).  ^[c]^Descriptors of defect density and defect type are the integrated intensity ratios: A_D_/A_G_, and A_D_/A_D'_ respectively. A_D_/A_D'_ of 13 and 7 are associated with sp^3^-defects and vacancy-like defects, respectively.  ^[d]^Area-weighted particle size distribution.  ^[e]^Analysed by RI and BF combined with the particle searching algorithm.  ^[f]^Determined from thermograms at T= 600 ˚C. | | | | | |

**Table S7.** Effect of GOs on the lifespan and health span of worms.

|  | **Vehicle** | **GO1** | **GO2** | **GO3** | **GO4** |
| --- | --- | --- | --- | --- | --- |
| Sample size | 50 | 50 | 50 | 50 | 50 |
| Censor | 11 | 8 | 18 | 32 | 17 |
| **Median of survival**  (days ± SEM) | 16.92 ± 0.66 | 15.16 ± 0.72* | 16.62 ± 0.61 | 15.97 ± 0.65* | 13.03 ± 1.06** |
| **Median of moving worms**  (days ± SEM) | 16.12 ± 0.69 | 15.19 ± 0.73 | 16.18 ± 0.69 | 14.74 ± 0.38 | 12.96 ± 1.05** |

Data are the mean ± SEM. * *p* <0.05 and ** *p* < 0.01 *vs*. vehicle according to log-rank and Bonferroni’s *post hoc* test between the pooled animal populations. The data are obtained from 50 worms from three independent experiments.

**Table S8.** LC_50_ values with 95% confidence intervals for *D. magna* exposed to GO particles for 48 hours.

| **GO** | **LC_50_ (mg/L)** | **CI 95%** |
| --- | --- | --- |
| GO1 | 118.9 | 114 – 123 |
| GO2 | 234.8 | 198 – 271 |
| GO3 | 140.9 | 122 – 159 |
| GO4 | 98.5 | 95 – 102 |
